# A Hedgehog-FGF signaling axis patterns anterior mesoderm during gastrulation

**DOI:** 10.1101/736322

**Authors:** Alexander Guzzetta, Mervenaz Koska, Megan Rowton, Junghun Kweon, Hunter Hidalgo, Heather Eckhart, Rebecca Back, Stephanie Lozano, Anne M. Moon, Anindita Basu, Michael Bressan, Sebastian Pott, Ivan P. Moskowitz

## Abstract

The application of single cell technologies to early development holds promise for resolving complex developmental phenotypes. Here we define a novel role for Hedgehog (Hh) signaling for the formation of anterior mesoderm lineages during gastrulation. Single-cell transcriptome analysis of Hh-deficient mesoderm revealed selective deficits in anterior mesoderm populations that later translate to physical defects to anterior embryonic structures including the first pharyngeal arch, heart, and anterior somites. We found that Hh-dependent anterior mesoderm defects were cell non-autonomous to Hh-signal reception. Transcriptional profiling of Hh-deficient mesoderm during gastrulation revealed disruptions to both transcriptional patterning of the mesoderm and a key FGF signaling pathway for mesoderm migration. FGF4 protein application was able to restore cellular migration during gastrulation that was decreased by Hh pathway antagonism. These findings implicate that primitive streak-mediated regulation of anterior mesoderm patterning is controlled by a multicomponent signaling hierarchy activated by Hh signaling and executed by FGF signal transduction.

**SIGNIFICANCE STATEMENT:** How signaling events during gastrulation pattern the mesoderm is a fascinating developmental process. Although Hedgehog signaling has been implicated in early mesoderm development, its mechanistic role has not been described. We applied single cell sequencing to describe mesodermal defects in Hedgehog pathway mutants—revealing selective defects in anterior mesoderm populations. Transcriptional profiling of gastrulating Hedgehog mutants indicated that several pathways essential for primitive streak function, including FGF, required Hh signaling. Blocking Hedgehog signaling abrogated cell migration during gastrulation, which could be mitigated by addition of FGF4 ligand. This work uncovers a novel Hedgehog to FGF signaling event and describes a unique mechanism by which signals from the node impact to anterior mesoderm formation through the modulation of primitive streak function.

## INTRODUCTION

One of the first challenges during metazoan development is to generate and distribute mesoderm across the anterior-posterior (A-P) axis during gastrulation (1–4). Presumptive mesoderm migrates through the primitive streak and is patterned during migration towards the anterior embryonic pole (1, 3–7). A-P patterning of the mesoderm is determined by secreted signals from the node and primitive streak. Midline-resident signaling pathways, including the Nodal, Fibroblast Growth Factor (FGF), and Wnt families, are all required for the patterning of anterior mesoderm (6, 8). While the requirement for these pathways in gastrulation have been defined by classical loss-of function assays, the multicomponent cross-talk between axis signaling mechanisms dictate A-P patterns remain largely unknown.

The Hedgehog (Hh) signaling pathway is required the morphogenesis of organs derived from all three germ layers in most metazoans (9, 10). Hh signaling was first described in a classic forward genetic screen for genes that determine A-P segment polarity during early *Drosophila melanogaster* development (11). Hh pathway activation in mammals is initiated by the binding of Hh ligands, *Shh*, *Ihh* or *Dhh*, to *Ptch1*—relieving its inhibition on Smoothened (*Smo*), which induces nuclear translocation of full-length Gli2/3 transcription factors (TFs) to promote Hh target gene transcription (12, 13). Conversely, the absence of Hh signaling triggers proteolysis and truncation of Gli2/3 proteins to their repressive isoforms Gli2R/3R (13, 14). Hh signaling works to pattern a diverse array of structures including the *Drosophila* wing disc (15), cnidarian pharyngeal musculature (16), tetrapod forelimb (17), vertebrate central nervous system (18) and heart (19–21). However, it has not been implicated in patterning the early A-P axis in vertebrates.

Removal of all Hh signaling through germline deletion of *Smo* in mice revealed an essential role for the Hh pathway in early mammalian development (22). *Smo^−/−^* embryos exhibit cardiac and somitic defects, absence of the anterior dorsal aorta, and fail to establish left-right (L-R) axis asymmetry (22, 23). Furthermore, treatment of zebrafish embryos with a small molecule *Smo* antagonist, cyclopamine, during early, but not late, gastrulation resulted in cardiac hypoplasia—suggesting that Hh signaling is essential during early embryogenesis for mesoderm lineage formation (24). Whereas *Shh* is the only Hh ligand necessary for L-R axis patterning in mammals (25), compound *Shh*^−/−^;*Ihh*^−/−^ mutants produce phenotypes indistinguishable from *Smo^−/−^* mutants (22). This suggests an important but poorly understood role for redundant *Shh* and *Ihh* signaling during early embryogenesis—independent of L-R patterning.

We investigated the role of Hh signaling for patterning the mesoderm during gastrulation. We observed that Hh signaling is first active, at E7.25, within the node, a critical midline structure that organizes L-R and A-P mesoderm patterning during gastrulation. We studied the consequence of early disruptions to midline Hh signaling in the mesoderm through single cell RNA-seq (scRNA-seq) by profiling wild type (Wt) and Hh-mutant mesoderm at the end of gastrulation. We identified a uniform deficiency in anterior mesoderm lineages in Hh-mutant embryos. Mesoderm-intrinsic and germline Hh pathway mutants both demonstrated later selective defects to anterior mesoderm-derived organs, including the head, heart, pharyngeal arch, and anterior but not posterior somites—consistent with a requirement for Hh signaling for anterior mesoderm development. Surprisingly, genetic inducible fate mapping showed that affected anterior mesoderm lineages did not appreciably receive Hh signaling. Transcriptional profiling of Hh-deficient mesoderm during gastrulation revealed disruptions to genes required for patterning at the primitive streak and the generation of anterior mesoderm, including the FGF signaling pathway. Small molecule Hh pathway inhibition in chick embryos caused dose-dependent mesoderm migration defects. Addition of FGF4 protein restored mesoderm migration following Hh inhibition. These findings resolve a node to primitive streak midline signaling axis, involving Hh and FGF signals respectively, responsible for the migration and patterning of anterior mesoderm during gastrulation.

## RESULTS

### Hedgehog signaling is first active in the organizing centers of embryonic axis determination

In order to study the role of early Hh pathway activity on mesoderm development, we set out to identify the earliest tissue to receive Hh signaling using a *Ptch1^LacZ^* reporter allele (26). No evidence of *Ptch1* reporter activity was observed prior to E7.0 (Fig. 1*A*) but becomes apparent in the node at E7.25, consistent with previous reports (Fig. 1*A*)(27). At E8.25, *Ptch1* reporter activity expands to the notochord (Fig. 1*A*) and shortly thereafter appears throughout the neural floorplate, somites, dorsal aorta and the SHF at E8.5 (Fig. 1*A*). These observations indicate that Hh signaling is first active in the node which is especially important for determining both L-R and A-P mesoderm patterning by mediating midline signals for mesoderm formation.

**Fig. 1.**
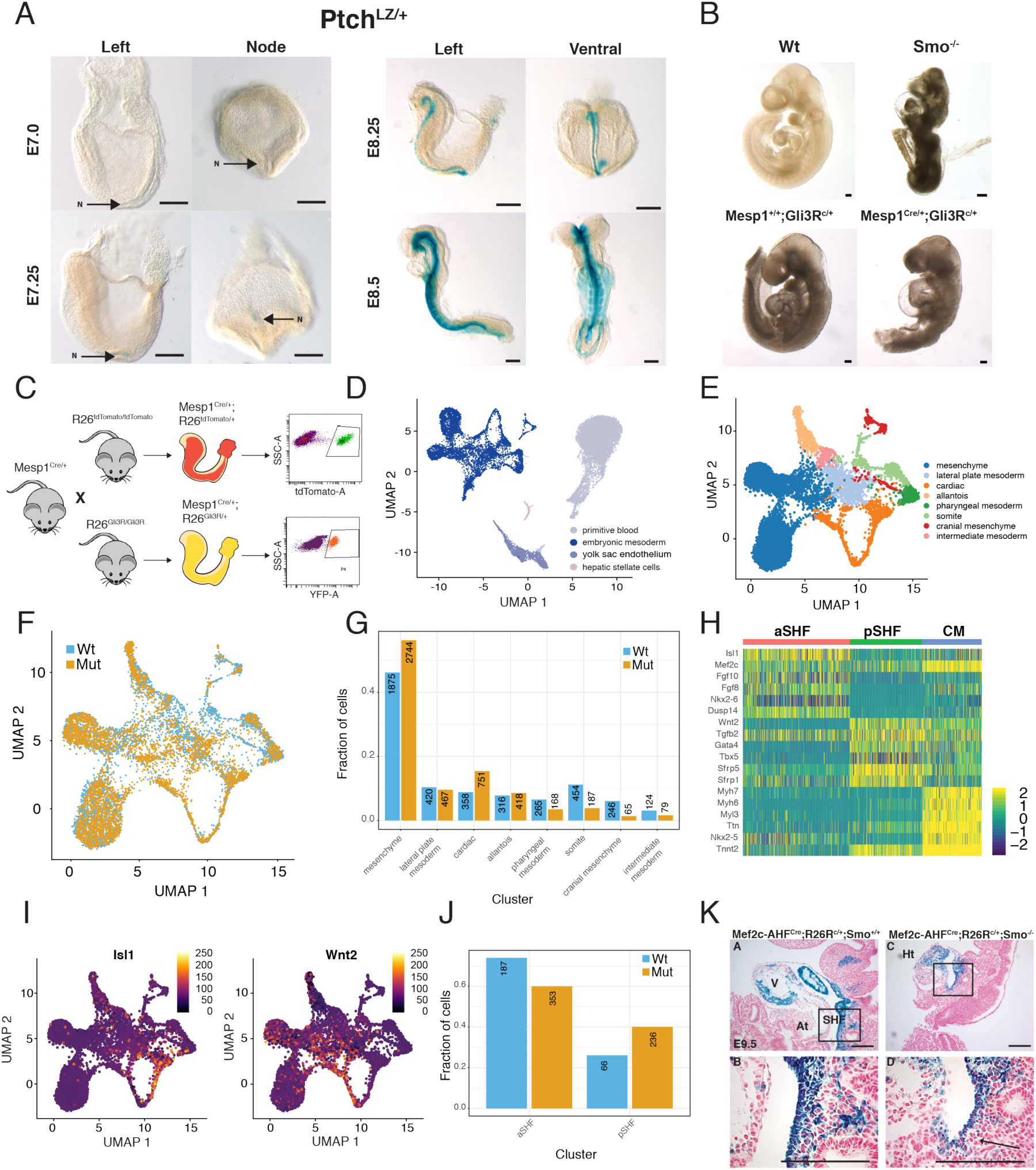
Single cell RNA-seq identifies Hh-dependent selective defects to anterior mesoderm progenitor populations. (A) Whole-mount views showing Ptch^Lz^ reporter activity which marks Hh receiving cells during early development between E7.0 (top-left) and E8.5 (bottom-right). (B) Whole-mount left lateral views of control and Hh-mutant embryos comparing defects observed between Smo^−/−^ mutants (top-right) and Mesp1^Cre/+^R26^Gli3R^ mutants (bottom-right). (C) Experimental schema for collecting mutant (R26-Gli3R) and wild type (R26-tdTomato) Mesp1^Cre^-labeled cells by FACS for single-cell RNA-seq. (D) Uniform manifold approximation and projection (UMAP) for all *R26-tdTomato and R26-Gli3R* cells used in this study. Extra embryonic mesoderm and blood are colored grey and light blue, while embryonic mesoderm lineages are shaded in dark blue. (E) UMAP representation of embryonic-derived mesoderm lineages where cells are colored by their cell type annotation with labels provided on the side of the plot. (F) UMAP for embryonic mesoderm with color segregation of control *R26-tdTomato* (blue) and mutant *R26-Gli3R* (orange) derived cells. (G) Column graph representing the proportion of *R26-tdTomato* (blue) *and R26-Gli3R* (orange) cells in each annotated lineage from (E) where the number linked to each column represents absolute cell counts. (H) Heatmap of marker gene expression for aSHF (red), pSHF (green), and cardiomyocytes (blue) where each column represents a single cell. Expression of aSHF and pSHF markers Isl1 and Wnt2 (I) superimposed on UMAP clusters derived from (E). (J) Proportion of *Mesp1^Cre^-tdTomato* (blue) *and Mesp1^Cre^-Gli3R* (orange) cells across aSHF, pSHF, and CM populations. (K) Fate map of aSHF-specific Cre (*Mef2c^AHFCre^*) with a Cre-dependent lacZ reporter (*R26R*) in *Smo*^+/+^ and *Smo*^−/−^ backgrounds. Images represent the posterior margin of the aSHF at low and high power in the top and bottom panels, respectively. (Legend: LV=left ventricle, SHF = Second Heart Field, At = Atrium, HT = Heart tube). Scale bars = 200 µm.

Based on the importance of signals sent from the node to mesoderm development, we assessed the necessity of Hh signaling for early mesoderm development by driving expression of a dominant-negative transcriptional repressor of the Hh pathway, *Gli3R* in the nascent mesoderm using Mesp1^Cre^ (28, 29). We observed severe head and heart tube defects in *Mesp1^Cre/+^;R26^Gli3R-IRES-Venus/+^* (*Mesp1^Cre^-Gli3R*) (30) mutants which phenocopied the majority of defects characteristic of Smo^−/−^ mutants (Fig. 1*B*). Both *Smo^−/−^* and *Mesp1^Cre^-Gli3R* mutants displayed abnormalities ranging far beyond L-R patterning defects alone and suggested an essential role for Hh signaling across early mesoderm development.

### Drop-seq reveals anterior mesoderm deficit in Hedgehog signaling mutants

To examine the consequence of disrupting mesoderm-intrinsic Hh signaling, we performed single-cell (sc) RNA-seq (Drop-seq) (31) on cells isolated by fluorescence activated cell sorting (FACS) from mutant (*Mesp1^Cre^-Gli3R*) and control *Mesp1*^Cre/+^;*R26*^tdTomato^ (*Mesp1^Cre^-tdTomato*) (32) embryos at the onset of organogenesis (E8.25)(33, 34)(Fig. 1*C*). We jointly processed and analyzed 9,843 *Mesp1^Cre^-tdTomato* and 10,663 *Mesp1^Cre^-Gli3R* cells from two biological replicates and observed similar gene detection across replicates (Fig. S1). Using unsupervised clustering to identify distinct cell populations (35, 36) we assigned cells to either embryonic mesoderm lineages (44%; 8,937/20,506) or hepatic stellate cells and extraembryonic tissues which are involved in early blood development (56%, 11,569/20,506) (Fig 1*D*, Fig. S2 and Table S1)(37). Focusing our analysis on the embryonic mesoderm, we identified all expected *Mesp1^Cre^*-derived lineages in both mutant and control embryos using a combination of canonical maker gene detection and correlation analysis with extant scRNA-seq datasets (Fig. S3, Fig. S4 and Fig. S5)(29, 38–40). When projected onto a Uniform Manifold Approximation and Projection (UMAP) plot, the proximity between clusters roughly recapitulated developmental relationships (Fig. 1*E*) with and without sample integration (41) (Fig. *1E* and Fig. *S6*). Specifically, we identified that the majority of embryonic mesoderm cells contributed to mesenchyme (4,619, 51.7%, dark blue) followed by: cardiac (1109, 12.4%, orange), lateral plate (887, 9.92%, grey-blue), allantoic (734, 8.21%, pale orange), somitic (641, 7.17%, light-green), pharyngeal (433, 4.85%, dark-green), cranial (311, 3.48%, red) and intermediate (203, 2.27%, salmon) mesoderm lineages (Fig. 1*E*, Fig. S3, and Fig. S4).

Expression of *Gli3R* significantly altered the distribution of *Mesp1^Cre^*-derived cells (Fig. 1*F*). We observed a selective deficiency in the proportional contribution of cells to anterior mesoderm lineages including cranial, pharyngeal and somitic mesoderm (Fig. 1*G*). Cranial mesoderm, marked by *Otx2* expression (42), is the anterior-most mesoderm lineage and demonstrated the most pronounced deficiency at more than four-fold reduction in lineage contribution from mutant *Mesp1-Gli3R* cells, compared to *Mesp1-tdTomato* controls (1.33% vs 6.06% of cells). Somitic mesoderm, which is marked by *Meox1* and *Tcf15* (43, 44) and contributes to the development of the anterior-most somites at E8.25, was reduced in mutants by nearly three-fold (3.82% vs. 11.2%). Finally, pharyngeal mesoderm, marked by *Col2a1* and *Sox9* (51, 52), demonstrated nearly 2-fold reduction within mutants (3.43% vs. 6.53%). Therefore, multiple independent anterior mesoderm lineages demonstrated a deficiency in *Mesp1^Cre^-* derived cell contributions after expression of the Hh-pathway trasncriptional repressor, *Gli3R*.

Interestingly the cardiac lineage, derived from the anterior mesoderm, was severely malformed in *Mesp1^Cre^-Gli3R* embryos—yet did not show the obvious proportional reductions in cell contribution. To investigate this further we analyzed the cardiac lineage in isolation and found it to be comprised of three distinct sub-clusters: functional cardiomyocytes marked by *Tnnt2* and *Myh6* (45, 46) and two cardiac progenitor clusters. Cardiac progenitors comprise a transient developmental structure called the second heart field (SHF) which is located anatomically dorsal to the heart tube (47). The SHF can be divided anatomically into anterior (aSHF) and posterior (pSHF) regions which express distinct transcripts. The aSHF is defined by *Fgf8, Fgf10* and *Isl1* expression (Fig. 1 *H* and *I*) (48) while the pSHF is defined by *Wnt2* and *Tbx5* expression (Fig. 1 *H* and *I*) (49, 50). Within the SHF populations mutant embryos contribute proportionally less cells to the aSHF (57.3%) compared to controls (73.9%) (Fig. 1*J*). To directly test whether Hh mutants exhibited decreased aSHF cellularity, we performed a genetic fate map using (*Mef2c^AHF-Cre^*), which expresses Cre recombinase primarily in the the aSHF (51). We analyzed the aSHF progenitor pool in *Smo*^+/+^ and *Smo*^−/−^ backgrounds using *Mef2c^AHF-Cre^* and a Cre-dependent lacZ reporter (*R26R*)(52). We observed a substantial reduction of the aSHF in Mef2c^AHF-Cre^;R26R^c/+^;*Smo*^−/−^ mutants compared to Mef2c^AHF-Cre^;R26R^c/+^;Smo^+/+^ controls at E9.5, when the aSHF is normally well established (Fig 1*J*). These data suggest that reduction in Hh signaling cause anterior-selective defects within the cardiac lineage, just as the embryo overall shows a deficiency of anterior lineages in *Mesp1-Gli3R embryos*.

### Hedgehog signaling is selectively required for anterior mesoderm development

Drop-seq analysis suggested that anterior mesoderm lineages were selectively reduced in Hh mutants during organogenesis. We hypothesized that this disruption would culminate in anterior-specific phenotypic defects later in development. Anterior embryonic mesoderm lineages arise during gastrulation when undifferentiated epiblast cells migrate through a transient structure known as the primitive streak (1, 3–5). The earliest cells to contribute to the embryo enter the streak during early to mid-streak stage and migrate furthest towards the anterior embryonic pole (4, 5) where they subsequently differentiate into specific lineages including, from anterior to posterior, pharyngeal mesoderm, cardiac mesoderm, and anterior somitic mesoderm (Fig. 2*A*). We directly analyzed the development of anterior mesoderm-derived structures in both mesoderm-intrinsic and germline Hh pathway mutants. Mesoderm-intrinsic Hh mutants, including Mesp1^Cre/+^;Smo^f/−^ (29, 53) *and Mesp1-Gli3R* embryos, exhibited cardiac and first pharyngeal arch hypoplasia compared to Mesp1^Cre/+^;Smo^f/+^ controls (Fig. 2*B* 2*C*). Although the anterior-most somites in mesoderm-intrinsic Hh mutants were positioned normally, they exhibited severe morphologic defects and failed to compact (Fig. 2*E* and 2*G*), in contrast to the anterior somites which were relatively similar to Mesp1^Cre/+^;Smo^f/+^ controls (Fig. 2*E* and 2*G*). Germline removal of *Smo* resulted in the most severe anterior defects including agenesis of the first pharyngeal arch and previously published cardiac defects (Fig 2*J*)(22). *Smo^−/−^* mutants also revealed a complete absence of the anterior-most somites, which were replaced by loosely packed mesenchyme (Fig. *2M*). Surprisingly, removal of *Shh* failed to phenocopy any aspect of the cardiac chamber or somite defects observed in *Smo^−/−^* mutants—instead, *Shh* mutants only exhibited previously described pharyngeal arch hypoplasia (Fig. 2*H*, 2*I*, 2*K* and 2*J*) (54). These results demonstrated that severely reduced Hh signaling by mutations to *Smo,* but not *Shh*, selectively affected the development of anterior mesoderm (22).

**Fig. 2:**
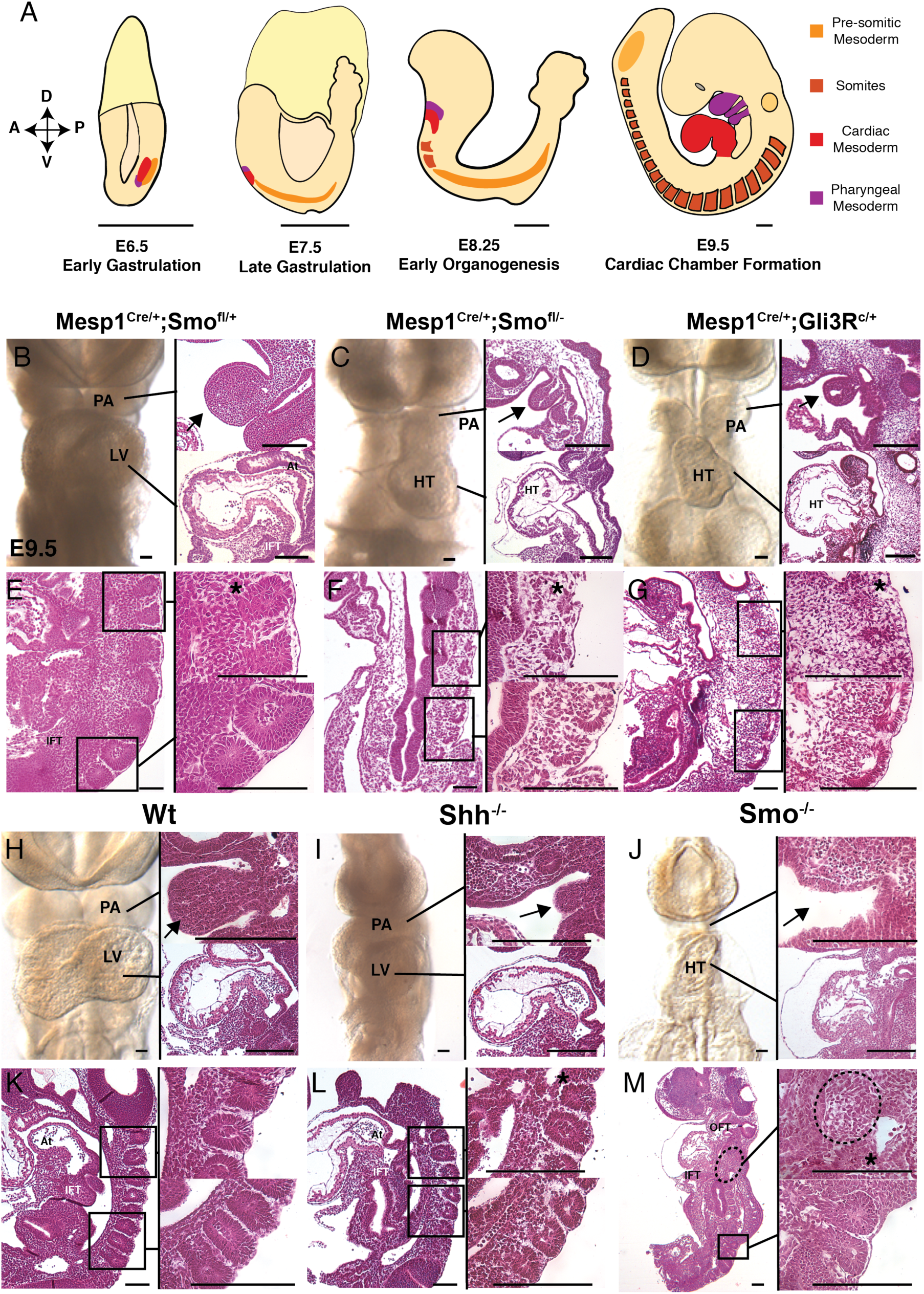
Hh signaling is required for anterior mesoderm morphogenesis. (*A*) Represents a cartoon model depicting the developmental ontogeny of anterior mesoderm lineages with estimated size bars for each embryonic stage. (*B*-*G*) demonstrate phenotypic abnormalities in conditional Hh mutants, *Mesp1^Cre/+^Smo^f/−^* (*C* and *F*) and Mesp1^Cre/+^;R26^Gli3R/+^ (*Mesp1-Gli3R)* (*D* and *G*) with their respective control, *Mesp1^Cre/+^;Smo^f/+^* (*B* and *E*) at E9.5. (*H*-*M*) Represent germline Hh pathway mutants, *Shh^−/−^* (*I* and *L*) and *Smo^−/−^* (*J* and *M*) and their respective Wt control (*H* and *K*) at E9.5. (*B*-*D* and *H*-*J*) Represent frontal whole-mount views of embryos (left panels) with corresponding sagittal histology sections of the pharyngeal arch (top-right panels) and heart (bottom-right panels). Black bars link the whole mount views of the pharyngeal arch and heart to their corresponding histology section. (*F*-*G* and *K*-*M*) Represent low-power sagittal histology views of both anterior and posterior somites (left panel). High power views of anterior and posterior somites are represented in the top-right panels and bottom-right panels, respectively. Somite 1 is indicated by an * in each top-right panel. (Scale bars = 200 µm). (Legend: IFT = Inflow tract; LV = Left Ventricle; OFT = Outflow tract; At = Atrium; PA = pharyngeal arch; D = Dorsal; A = Anterior; P = Posterior; V = Ventral).

### Hh reception by cardiac progenitors is not required for early heart development

Due to the selective role for Hh signaling in anterior-specific mesoderm development, we hypothesized that anterior lineages would autonomously require intact Hh pathway function. To test this in the cardiovascular lineage, we conditionally removed *Smo* from the earliest specified cardiac precursors using *Nkx2.5^Cre^* (55). Surprisingly, no phenotypic abnormality was observed in *Nkx2.5^Cre/+^;Smo^f/−^* embryos (Fig. S7*B*) compared to *Nkx2.5^Cre/+^;Smo^f/−^* controls (Fig. S7*A*) at E9.5. To rule-out an insufficiency of Nkx2.5^Cre^ to functionally delete *Smo* in cardiac progenitors we analyzed *Nkx2.5^Cre/+^;Smo^f/−^* embryos later in development where Hh signaling is required within the *Nkx2.5* sub-domain of the SHF for atrioventricular septum development (21, 56). At E14.5, *Nkx2.5^Cre/+^;Smo^f/−^* displayed completely penetrant atrioventricular septal defects (AVSDs) (Fig. S7D) compared to normal morphology observed in *Nkx2.5^Cre/+^;Smo^f/+^* littermate controls (Fig. S7*C*). Additionally, genetic fate mapping of *Nkx2.5^Cre^* expression revealed no Hh dependence in that we observed identical fate maps between *Nkx2.5^Cre/+^;Smo^f/−^;R26R^c/^*^+^ mutants and *Nkx2.5^Cre/+^;Smo^f/−^;R26R^c/+^* controls where both genotypes were stained throughout the cardiac chambers and SHF (Fig. S7*E* and *F*). These data suggested that Hh signaling was either not cell-autonomously required for cardiac development or required for cardiac development in early pre-cardiac mesoderm prior to specification.

### Hh-dependent anterior mesoderm lineages do not directly receive Hh signaling

To ask whether anterior mesoderm lineages receive Hh signaling at any point during early development, we utilized an inducible CreERT2 knock-in allele at the *Gli1* locus (*Gli1*^CreERT2^)(57) to mark Hh-receiving cells and their decedents in a tamoxifen (TM)-dependent manner. TM was administered between E5.5 and E8.5 in *Gli1^Cre/+^;R26R^c/c^* embryos and analyzed at E9.5 (Fig. 3 *A*). We observed a near-absence of marked cells when TM was administered one day preceding gastrulation at E5.5 (Fig 3*A*)—suggesting that Hh signaling is not active in embryonic mesoderm immediately prior to gastrulation. Administration of TM during early gastrulation at either E6.5, or late gastrulation at E7.5, resulted in robust X-gal staining throughout the neural tube and lateral mesenchyme; notably, few to no positive clones appeared in the pharyngeal mesoderm, somite bodies, or heart (Fig 3*A*). TM administered shortly after gastrulation at E8.5 demonstrated much more prevalent staining in the ventral neural tube and paraxial mesoderm but anterior mesoderm lineages remained unmarked (Fig. 3*A*). TM administration at both E6.5 and E7.5 did not increase the number of marked lineages, although we did observe higher proportion of stained cells within previously marked lineages (Fig 3*B*). Serially sectioned embryos revealed a near-absence of labeled clones in the pharyngeal mesoderm and somite bodies (Fig 3*C*). In contrast to the heart itself, which had few to no labeled clones throughout several litters analyzed (Table *S*1, Fig 3*D*), SHF cells were robustly labeled—providing a positive control for a lineage with known Hh signaling activity (Fig. 3*D*)(21, 56, 58). Collectively, the striking absence of Hh-activated cells within anterior organ rudiments suggested that while Hh is required for proper patterning of somatic, pharyngeal, and cardiac structures, this process is likely not mediated by direct signaling events.

**Fig. 3:**
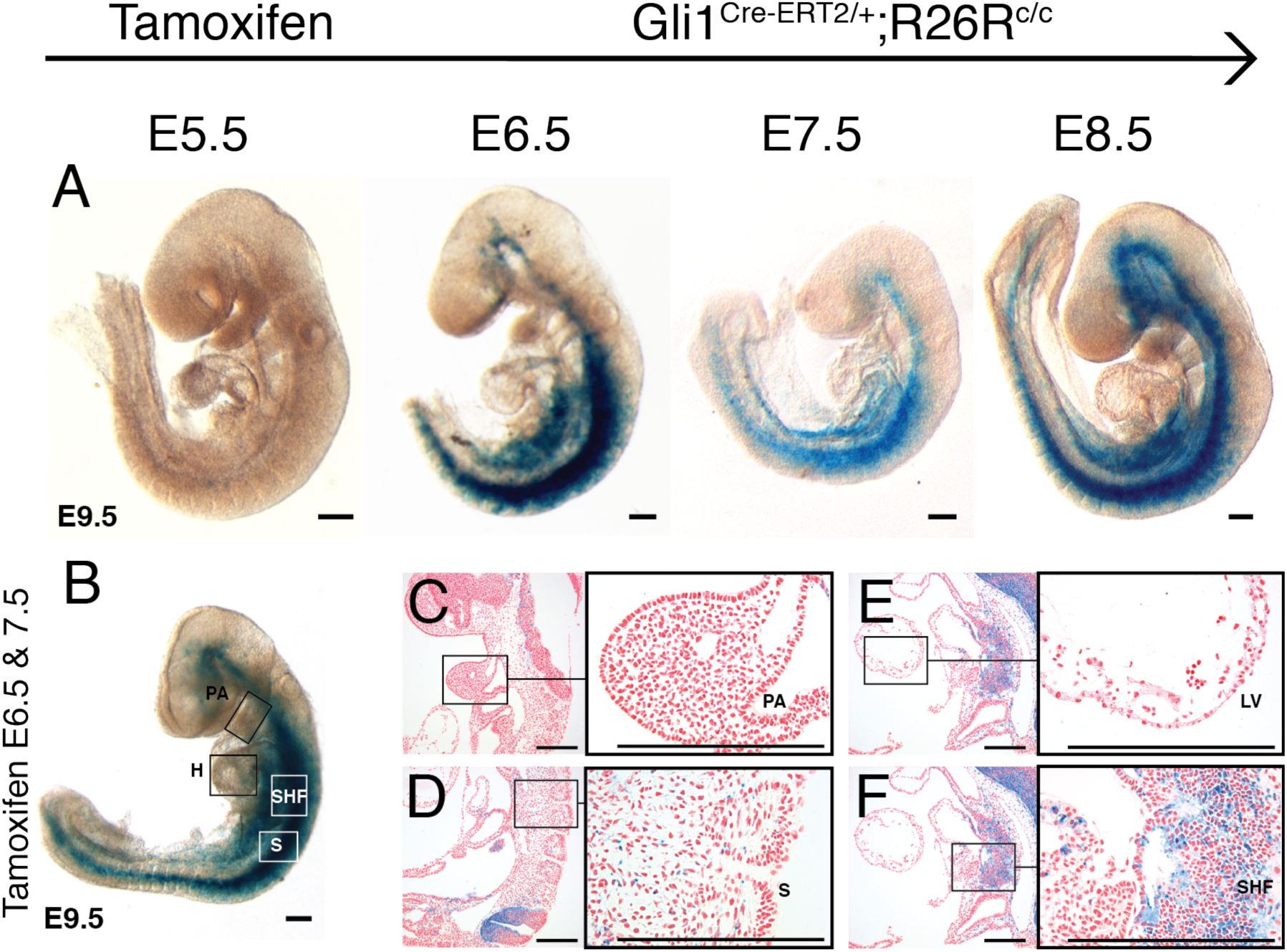
Anterior mesoderm lineages do not receive Hh signaling during gastrulation. (*A*) Shows genetic-inducible fate maps for Hh-receiving cells in Gli1^CreERT2/+^; R26R^c/c^ embryos from whole-mount left lateral views at E9.5. Embryos were harvested from pregnant dams given a single tamoxifen dose at the times indicated at top. (*B*-*F*) Shows whole mount *(B)* and sagittal sections (*C-F*) from embryos harvested from pregnant dams dosed with tamoxifen at both E6.5 and E7.5. (*C*) Shows sagittal sections of the pharyngeal arch (*C*), anterior somites (D), left ventricle (*E*), and second heart field (*F*) at low (left) and high(right) magnifications respectively; black boxes in the low-magnification field are linked to the area of high magnification. (Size bars = 200um) (Legend: PA = Pharyngeal arch; H = heart; SHF = Second Heart Field; S = somite).

### Transcriptional profiling identified Hh-dependent transcription factors and signaling pathways necessary for mesoderm morphogenesis

To identify candidate Hh-dependent genes capable of initiating anterior lineage development through a secondary signaling event, we performed RNA-seq on *Mesp1^Cre^-Gli3R* mutants late during gastrulation at E7.5. We utilized a Cre-dependent dual color system to separate red fluorophore-expressing wild type (*Mesp1^Cre^-Tomato*), from yellow fluorophore-expressing Hh mutant (*Mesp1^Cre^-Gli3R*) *Mesp1*^Cre^-labeled cells by FACS in order to generate three biological replicates of litter-matched RNA-seq samples (Fig. 4*A*). We observed consistent differential expression between biological replicates and identified 190 dysregulated genes by mRNA-seq (FDR <= 0.10) and (Fig. 4*B*). A large proportion of downregulated genes are known to be critical for primitive streak function and anterior mesoderm morphogenesis—including *Wnt3a*, *Msgn1*, Crypto (*Tdgf1*)*, Fgf4*, *Fgf8, and Smad3* (59–63). Additionally, classic factors for both A-P patterning and midline development, Brachyury (*T*) and Hnf3β (*FoxA2*), were also downregulated (64–66).

**Fig. 4:**
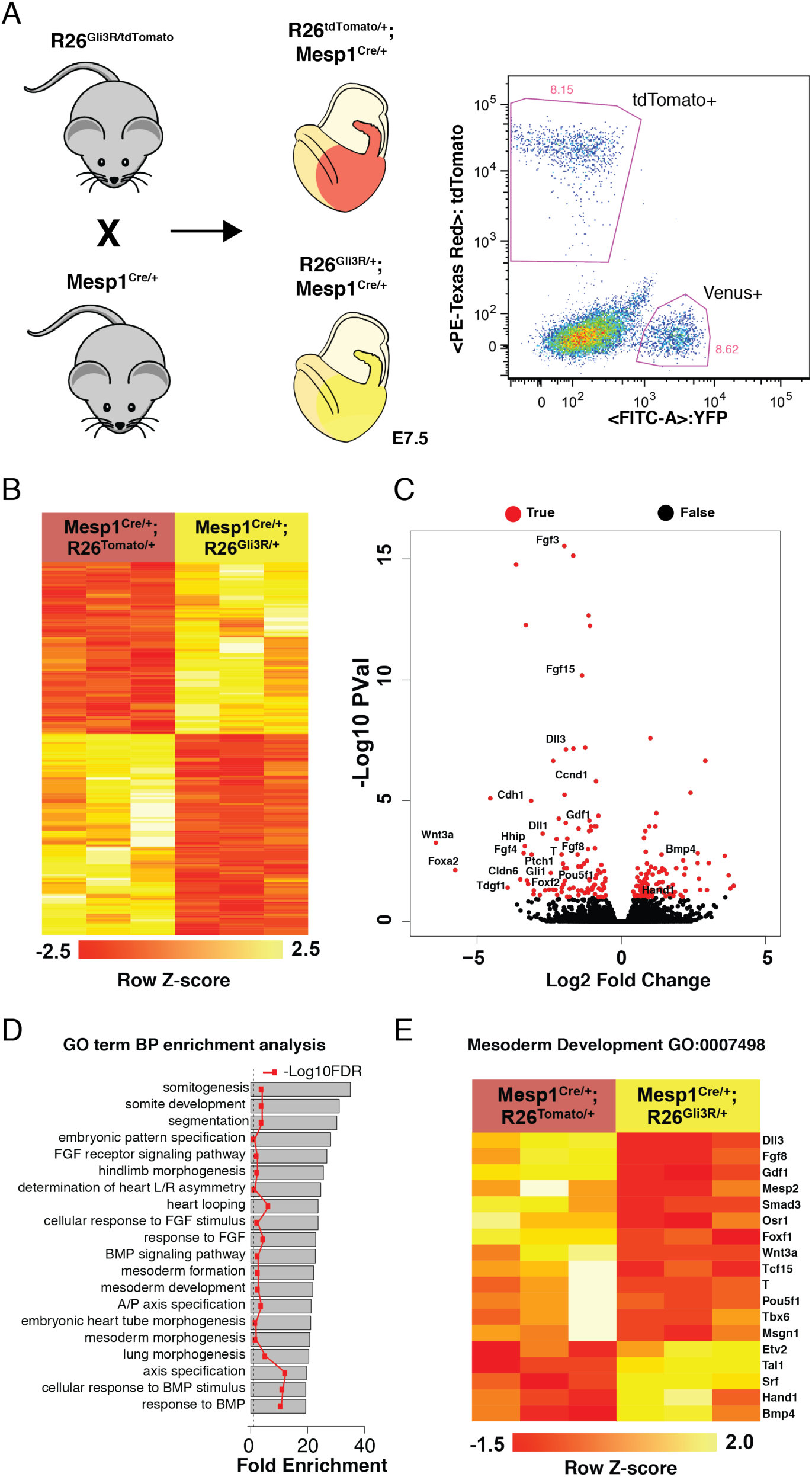
Disruption of Hh signaling causes major disruptions in genetic pathways for mesoderm development. (*A*) Represents the breeding strategy to produce litter-matched controls of Hh mutant (*Mesp1^Cre^-Gli3R*) and wild type (*Mesp1^Cre^-tdTomato*) mesoderm by FACS-sorting Mesp1^Cre^-marked cells by yellow and red channels respectively. (*B*) **Heat maps are labeled with row** Z-score normalized heatmap for dysregulated genes (FDR <= 0.10) where each column represents individual embryos. (*C*) A volcano plot for dysregulated genes in Hh mutants. (*D*) GO analysis for genes downregulated genes in *Mesp1^Cre^-Gli3R* mutants. (*E*) Heatmaps for all genes dysregulated under the Mesoderm Development (GO:007498) term (FDR <= 0.30).

Gene Ontology (GO) analysis for downregulated genes (FDR < 0.10) (Fig. 4*D*) confirmed our initial assessment that mesoderm morphogenesis was perturbed as nearly all top terms were associated with early mesoderm development—including the general term ‘Mesoderm Development’ (GO:0007498) (Fig. 4 *D* and *E*). The majority of significantly dysregulated genes (FDR <= 0.30) classified under ‘Mesoderm Development’ were downregulated— suggesting widespread dysfunction to primitive streak function and mesoderm morphogenesis. Interestingly, the few upregulated genes under ‘Mesoderm Morphogenesis’ such as *Bmp4 and Hand1,* are primarily involved in the development of the extraembryonic tissues at this stage (67, 68), which suggests a divergent role for Hh signaling between embryonic and extraembryonic tissues.

### Hh-dependent TFs are enriched for determinants of mesoderm morphogenesis and somitogenesis

GO analysis for Hh-dependent TFs identified somitogenesis and paraxial mesoderm, which share common genes for anterior mesoderm development during gastrulation, as the most highly represented terms (Fig. 5*A*). Genes disrupted within the GO term for somitogenesis (GO:0001756)(FDR <0.3) reveal downregulation of transcription factors that are important to both somite development alone (*Hes7*, *Tcf15*, *Dll3*, *Ripply2*)(69–73) and factors that also play a dual role in somite development and gastrulation (*Msgn1*, *Tbx6*, *Mesp2*, *FoxA2*, *T*, and *Smad3*)(59, 64–66)(Fig. 5*B*). Using RNA *in situ* hybridization, we analyzed the spatial expression patterns for key genes responsible for shared paraxial and mesoderm development (*Tbx6*, *Msgn1*, and *Mesp2*) along with Dll3, a gene responsible primarily for somitogenesis. *Tbx6*, which is critical for L/R patterning and mesoderm formation (59, 74), is lost from all but the most distal mesoderm expression domains in *Mesp1^Cre^-Gli3R* mutants (Fig. 5*C*). *Msgn1*, which is a regulates both pre-somitic and general mesoderm development (59, 75), lost expression in the majority of paraxial mesoderm and only maintained medial posterior expression—proximal to the primitive streak (Fig. 5*C*). *Mesp2,* which is crucial for both nascent mesoderm development and somite-pre-somitic mesoderm boundary determination (76), prematurely lost expression in the posterior streak while maintaining nascent somite boundary expression. Conversely, *Dll3* only appeared to lose a small portion of right-sided expression distal to the streak (Fig. 5*C*). In summary, all three shared mesoderm-somite program genes, but not the somitogenesis-only gene *Dll3*, demonstrated significantly reduced expression near the primitive streak in *Mesp1^Cre^-Gli3R* mutants (Fig. 5*C*).

**Fig. 5:**
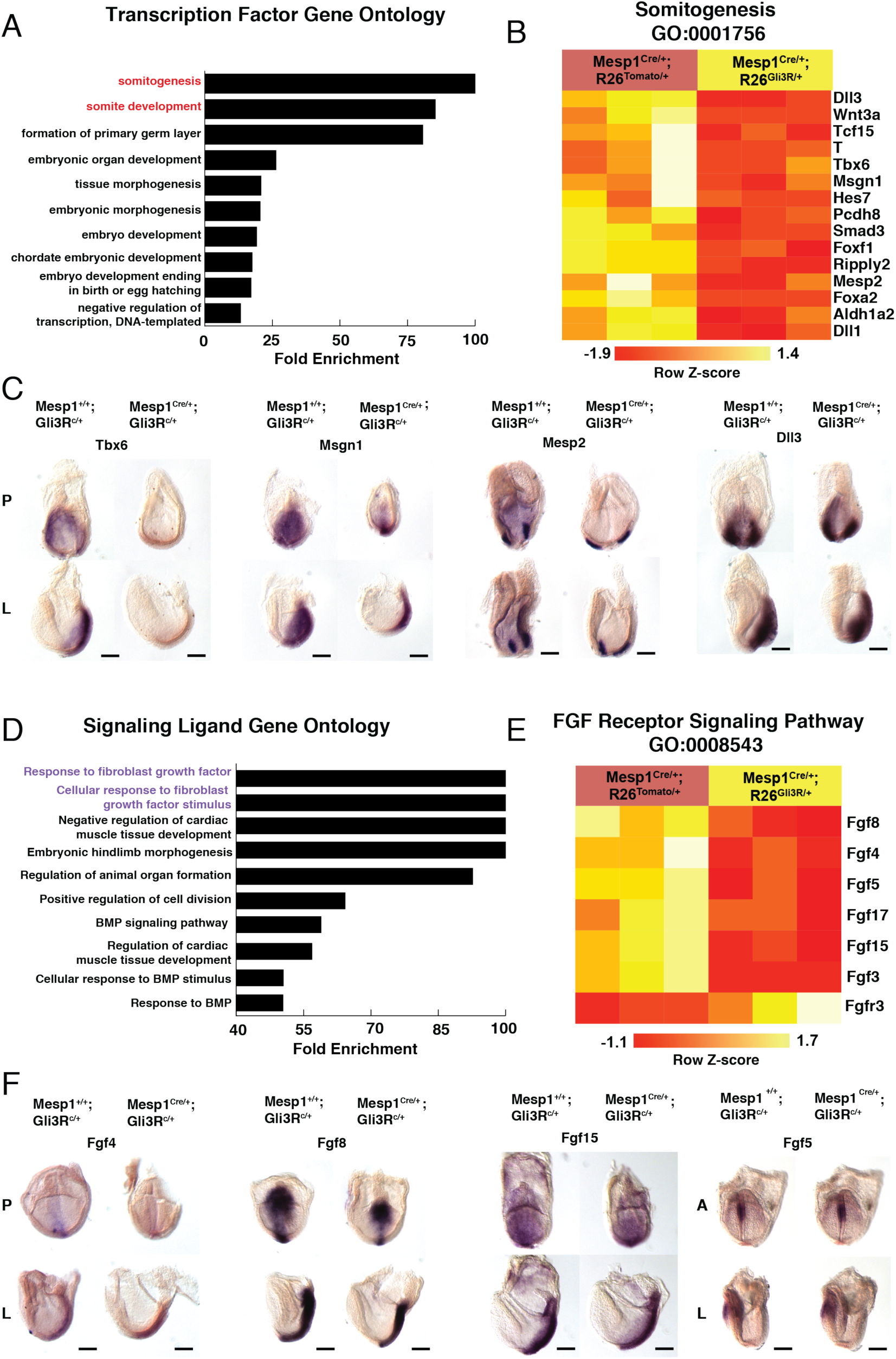
Targeted pathway analysis reveals widespread dysfunction of nascent mesoderm and Fgf pathways. (*A*) GO analysis performed only on downregulated genes classified as transcription factors (*B*) shows a heatmap generated from dysregulated genes (FDR <=0.30) overlapping with top GO term, Somitogenesis (GO:0001756). (*C*) ISH for the somitogenesis genes *Msgn1*, *Tbx6*, *Mesp2,* and *Dll1* in control (Mesp1^+/+^R26^Gli3R/+^) and Mut (Mesp1^Cre/+^;R26^Gli3R/+^) embryos at E7.5. (*D*) GO analysis for downregulated genes classified as signaling molecules. (*E*) Represents a heatmap for dysregulated genes FDR (<= 0.30) for the top GO term, FGF Receptor Signaling Pathway (GO:0008543. (*F*) ISH for FGF ligands *Fgf4, Fgf8, Fgf15,* and *Fgf5* in control (Mesp1^+/+^R26^Gli3R/+^) and Mut (Mesp1^Cre/+^;R26^Gli3R/+^) embryos at E7.5. (Scale bars = 200um) (Legend: A = Anterior, P = Posterior, L = Left).

### Transcriptional profiling identified FGF signaling downstream of Hh signaling for anterior mesoderm morphogenesis

We hypothesized that a secondary signaling pathway downstream of Hh signaling was required for anterior mesoderm development, based on the observation that anterior mesoderm lineages required intact Hh signaling despite not receiving Hh signaling directly. To identify candidate pathways, we performed GO analysis for downregulated genes classified as either signal ligands or ligand receptors in the Fantom consortium database (77) (Fig. 5*D*). The FGF pathway comprised the top candidate using this approach and differential expression of genes in the GO category “FGF Receptor signaling pathway” (GO:0008543) revealed a striking pattern of downregulation among all included FGF ligands (FDR <= 0.30) (Fig. 5*E*). Specifically, *Fgf4* and *Fgf8* share posterior expression domains in the primitive streak and both lose much of their posterior-lateral expression in Hh mutants (Fig 5*F*). Studies in early mammalian development have shown that the FGF pathway directs the migration of nascent mesoderm towards the anterior embryonic pole (61, 78) and *Fgf4* and *Fgf8* are uniquely important for this process (60, 61, 79). Given the quantitative and qualitative reductions to *Fgf4/8* expression in the primitive streak, we hypothesized that Hh signaling lies upstream of an FGF pathway for mesoderm migration.

### Gastrulation cell migration defects caused by Hh pathway antagonism are rescued by FGF4

To directly assess whether impaired Hh signaling disrupts mesoderm migration, we utilized a chick embryonic model of gastrulation. Mesoderm migration was tracked by adding either DiI (red) and DiO (green) fluorescent lipophilic dyes on each side of the primitive streak in Hamburger-Hamilton (80) stage 3 chick embryos (HH3) (Fig. 6*A*). Embryos treated with DMSO vehicle demonstrated classic anterior-lateral movement of labeled cells consistent with well-established fate and migration maps of nascent mesoderm (Fig. 6*B*) (61, 81), which could be quantified (Fig. 6*C*). We observed a dose-dependent reduction of nascent mesoderm migration to cyclopamine, a Hh pathway antagonist (82) (Fig. 6*F*). Embryos treated with 25 µM cyclopamine demonstrated a significant reduction in directional migration (ttest pValue < 0.005) (Fig. 6*D*) whereas embryos treated with 50 µM cyclopamine or higher demonstrated migratory defects compounded with widespread phenotypic disruptions including greatly reduced embryo size (Fig. 6*E*). These experiments demonstrated that A-P cell migration during gastrulation was hindered by Hh pathway inhibition (Fig. 6*F*).

**Fig. 6.**
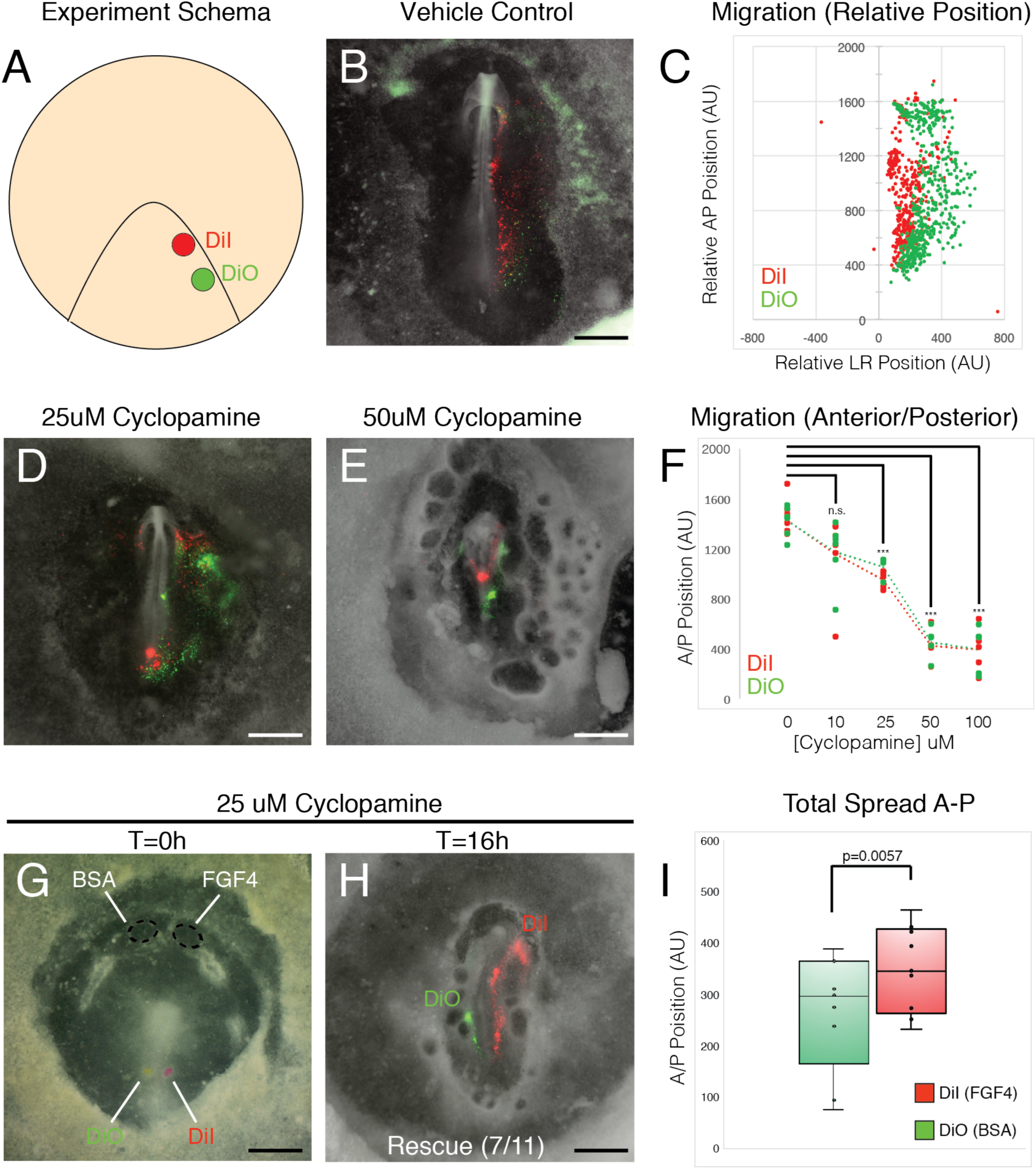
Hh signaling is required for anterior cell migration during gastrulation upstream of the FGF pathway. (*A*) Demonstrates the placement of two lipophilic dyes at the posterior portion of a HH stage 3 chick embryo (dorsal view). (*B and C*) Demonstrates the trajectory of post-ingression mesoderm cells in vehicle control-treated chick embryos (*B*) which is spatially quantified in (*C*). (*D and E*) Addition of Smo antagonist Cyclopamine at 25 µM (*D*) and 50 µM (E). (*F*) Shows quantification of inhibitory effect of cyclopamine between 10 µM and 100 µM. (*G-I*) embryos treated with 25 µM of cyclopamine had heparin beads coated with FGF4 placed towards the anterior-right pole of the embryo with BSA-coated heparin beads placed on the anterior-left portion of the embryo to serve as a contralateral control (*G*). After 16 hours of gastrulation 7/11 embryos show a marked rescue of specific directional migration on the FGF4-treated side (*H*) which was quantified to significantly differ from the control p=0.0057 (*I*). (Scale bars = 1mm).

To test whether FGF-directed cell migration was downstream of the Hh pathway, we placed beads coated with either a BSA control or FGF4 protein or a BSA control on opposite sides of the anterior midline in 25 µM-cyclopamine-treated HH4-stage chick embryos (Fig. 6*G*). We determined and quantified the effect of FGF4 on migration by monitoring the anterior excursion of DiO or DiI dyes of the primitive streak over a 16 hr period of gastrulation. Importantly, labeled cells that gastrulated on the same side as FGF4-coated beads showed a statistically significant increase in anterior-posterior spread when compared with cells migrating through the contralateral side of the primitive streak (towards control beads coated with BSA) (N=11; ttest pValue=0.0057; Fig. 6*H* and *I*). These data indicate that the migratory defects resulting from cyclopamine treatment can be mitigated by FGF4. These experiments provide evidence that Hh signaling is required upstream of FGF signaling for anterior mesoderm morphogenetic movements through the primitive streak during gastrulation.

## DISCUSSION

The mechanism by which embryos pattern tissues across their axes has fascinated developmental biologists since the founding of embryology. The application of single-cell sequencing can begin to unravel complex patterning defects by interrogating distinct cell types separately in a single assay. Leveraging this resource, we uncovered a novel role for Hh signaling in the development of anterior mesoderm lineages during gastrulation. We identified the dependence of nodal and FGF, critical signaling pathways for gastrulation, on intact Hh signaling. Finally, we demonstrated that FGF expression in the streak is downstream of the Hh pathway and that exogenous FGF replacement can rescue Hh-dependent migratory defects during gastrulation. Together these observations indicate that Hh signaling, observed only in the embryonic node during gastrulation, is essential for inducing migratory FGF signals from the primitive streak required for the appropriate allocation of mesoderm to developing anterior organs including the head, heart, pharynx and anterior somites. This work supports the application of single cell sequencing for the deconvolution of complex developmental phenotypes.

Although the FGF pathway is a well-known mitogen, its primary role during gastrulation is to direct coordinated cell migration through matrix remodeling and chemotaxis—particularly through the action of FGF4 and FGF8 (61). *Fgf8* germline hypomorph mutants reveal anterior-specific defects across multiple organ systems—including hypoplasia of the pharyngeal mesoderm, cardiovascular lineages, and selective anterior truncation of the zygomatic arch (83, 84). *Fgf4/8* conditional deletions within the late-streak posterior mesoderm reveal widespread somite defects distal to the posterior source of *Fgf4/8* expression (60, 79). The anterior-specific defects we observed in mesoderm-intrinsic Hh mutants phenocopy perturbations to FGF signaling in the streak where tissues distal to the signaling event are either absent or reduced. Our data provides clear evidence that one mechanism by which Hh signaling drives anterior mesoderm development is through the modulation of downstream *Fgf4/8* signaling. However, additional studies should address whether Hh signaling lies directly upstream of FGF pathway genes or works through an intermediary signal to affect FGF expression in the streak.

A prime candidate to serve as an intermediate between the Hh and FGF signaling is the Nodal pathway, which is active in the embryonic node concomitant with Hh pathway activity. Not only do Nodal pathway mutants exhibit anterior-specific defects during early development (6), there is evidence for cross-activation with the Hh pathway (22, 85–87). Although *Nodal* ligand is not disrupted in our mutants, we did observe Hh-dependence for two critical genes for Nodal pathway activity, *Tdgf1*, and *Smad3*. Furthermore, we observed a critical network of shared target genes downstream to both pathways, including *FoxA2*, *T*, and *Fgf4*—all of which are critical to A-P mesoderm patterning (63–66, 88). Specifically, the direct regulation of *Fgf4* expression and the indirect regulation of *Fgf8* through *T* provide strong circumstantial evidence for a role of Nodal to help drive FGF signaling in the primitive streak. This shared gene set suggests that the Hh and Nodal pathway may work in concert to modulate FGF gene expression in the primitive streak.

We demonstrated that redundant Hh signals in the early embryo serve a previously uncharacterized role for Hh signaling in the development of anterior mesoderm lineages. We note that cardiac, cranial, pharyngeal, and anterior somitic mesoderm migrate through the primitive streak at nearly the same time as Hh pathway activity arises within the embryonic node (7). Here, we provide a mechanistic link between Hh signaling from the node and anterior mesoderm development by establishing the Hh-dependence of the FGF pathway for cell migration during gastrulation (Fig. 7*A*). Perturbations to early Hh signaling diminish the expression of migratory FGF signals from the midline—leading to A-P patterning defects through dysfunctional mesoderm migration at the primitive streak (Fig. 7*B*). These findings are the first to provide a functional link between midline signals from the embryonic node and primitive streak to control the patterning of anterior embryonic organs, and may describe a general mechanism for mammalian A-P patterning during gastrulation.

**Fig. 7:**
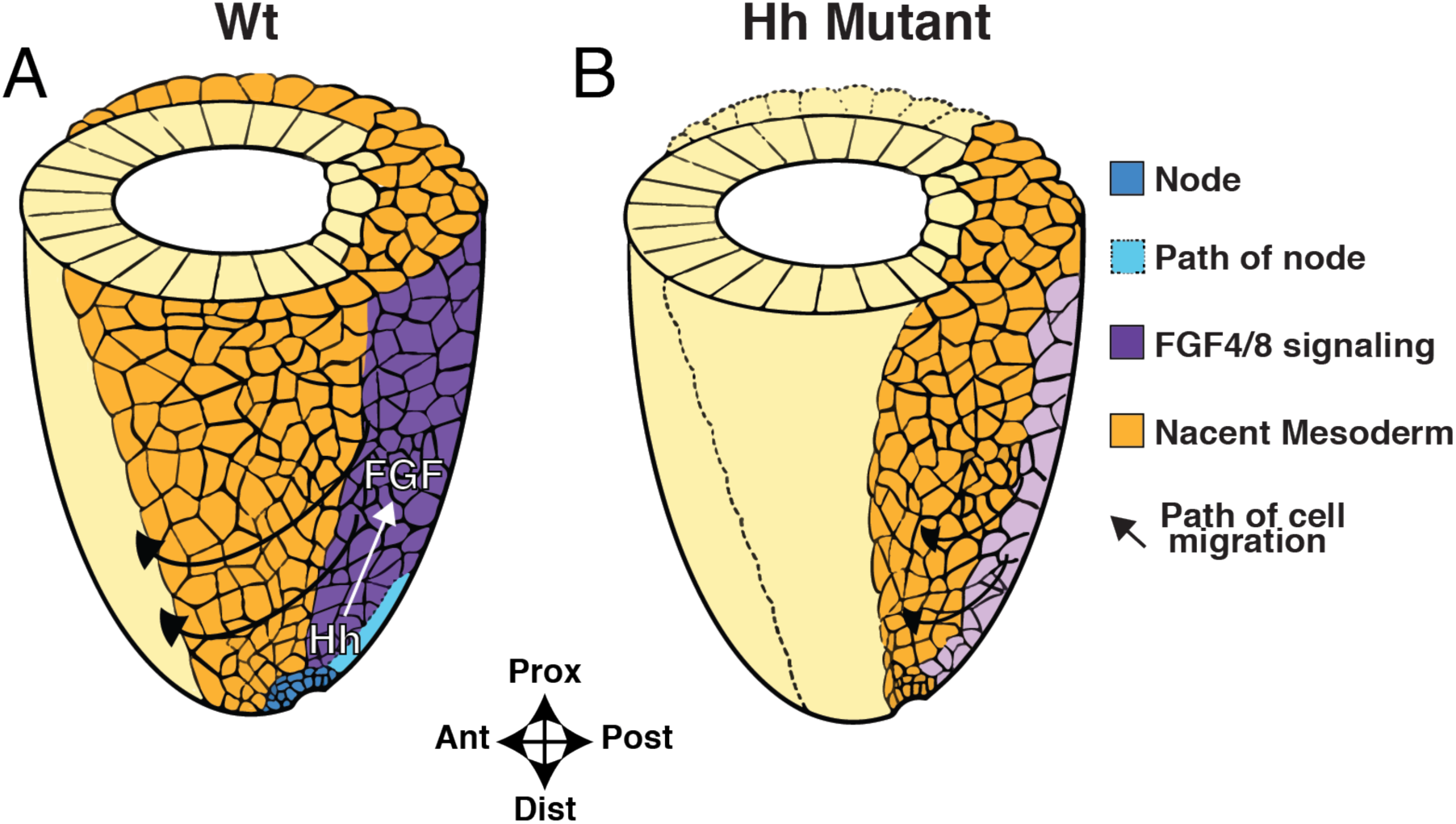
Midline Hh signaling from the node drives an FGF pathway for anterior mesoderm morphogenesis. (*A* and *B*) Demonstrate a cartoon model for the role of Hh signaling in anterior mesoderm patterning. (*A*) In Wt embryos, Hh signaling is active early in embryonic midline in the node which is required for the full activation of a midline *Fgf4/8* signaling axis from the primitive streak which drives mesoderm migration. (B) in Hh pathway mutants, the FGF migratory pathway is greatly attenuated—resulting in disproportionate impacts on the migration and patterning of anterior mesoderm lineages which must migrate furthest during development. (Legend: Prox = Proximal, Dist = Distal, Ant = Anterior, Post = Posterior).

## METHODS

### Animal husbandry genotyping

All animals were housed in a barrier facility at the University of Chicago Animal Resource Center. Mice were genotyped by PCR using genomic DNA extracted by incubating tail clips or embryonic tissue in a 50mM NaOH solution at 98°C for 45 minutes; prior to genotyping, crude DNA extracts were neutralized with a 10mM Tris-HCL pH 8 solution. Genotyping protocols were based on the conditions the Jackson Laboratory provided corresponding to each locus, including primer sequences and thermocycler conditions.

### Superovulation and tamoxifen administration

For large-scale mouse timed pregnancies 21-35 days old female mice were injected intraperitoneally (IP) with 5 units of pregnant mares serum (PMS) (EMD Chemicals) in 0.9% saline between noon and 4 PM. 46-48 hours after the PMS injection, mice received a second IP injection with 5 units of human chorionic gonadotropin (HCG) (Sigma). Immediately after injection females were placed into the cages of stud males, 8 weeks of age or older. The noon of the day a vaginal plug was observed was designated E0.5. For all other embryo collections, female mice of 6-16 weeks were housed with stud males of 8 weeks or older until the detection of a vaginal plug at noon the next day (E0.5). For lineage tracing experiments, mice were treated with 2 mg of tamoxifen (MP Biomedicals) mixed in a 2:1 ratio with progesterone by (Sigma) intraperitoneal injection.

### Mouse euthanasia and embryo collections

Embryo were harvested from pregnant dams after euthanasia by CO_2_ asphyxiation and cervical dislocation. Embryos between E7.5 to E14.5 of age were dissected in cold phosphate-buffered saline (1x PBS). In all experiments save for live-cell assays and LacZ staining, embryos were incubated in 4% paraformaldehyde (4% PFA):PBST overnight. Embryos used for RNA *in situ* hybridization were further dehydrated by one wash with 50% Methanol (MeOH):PBST and two subsequent washes in 100% MeOH before storage at −20°C.

### LacZ staining

Embryos were fixed 4% PFA:PBST for 20 minutes with gentle rocking at 4°C. Embryos were washed 3 times for 5 minutes in 1X PBS. They were then incubated overnight at 37°C with staining solution comprised of 5ml of 1x PBS, 1.5ml of 5M NaCl, 50µl of 1M MgCl2, 1.5 ml of 10% TX-100, 500 µl of 0.5M Potassium ferricyanide, 500 µl of 0.5M Potassium ferrocyanide in 50 ml of total volume. X-gal solution (100mg/ml in DMF) (RPI Corp) stored in −20°C in light-impermeable containers was added at a concentration of 1:100 immediately prior to staining. Following staining, embryos were washed three times with 1x PBS and re-fixed in 4% PFA for one hour at room temperature.

### Riboprobe design

Probe templates were amplified using primers for partial cDNA sequences. Transcript mRNA templates were taken from the NCBI refseq. Primers were designed to amplify partial cDNAs between 400-1000 base pairs long, with 45-55% GC content. On the 5’ end of the downstream (antisense) primer, the sequence 5’-CTAATACGACTCACTATAGGGAGA-3’ was added. This served as the minimal promoter site for T7 to increase the efficiency of the transcription reaction. The PCR product was ran on a 0.8% TAE gel and amplicons of the correct predicted sizes were isolated using QIAquick gel extraction kit (Qiagen). For the genes that PCR primers failed to reliably amplify off of cDNA templates, we utilized the commercially available Gibson assembly service (gBlocks) provided by Integrated DNA Technologies. 440 base pairs from mRNA sequences were chosen to design the gBlock, with paired gateway sites comprised of an Attb1 (GGGGACAAGTTTGTACAAAAAAGCAGGC) site upstream and Attb2-(GACCCAGCTTTCTTGTACAAAGTGGTCCCC) site downstream of the gBlock fragment. gBlock fragments were cloned into pDonor using BP clonase (Invitrogen). The reaction components were incubated 3 hours at room temperature. 1µl of ProK was added and the mix was incubated for 10 min at 37°C to stop the reaction. Gateway clones were then transformed into competent dH5α bacteria (Invitrogen). Plasmids were then isolated from individual bacterial colonies and validated for correct insertions using Sanger sequencing performed at the University of Chicago Sequence Facility.

### In vitro transcription

*In vitro* transcription of dioxigenin-labeled riboprobe was performed with 1 μg linearized plasmid template or 200 ng PCR template. The in vitro transcription mixture comprised of DNA template, 2 μl 10x transcription buffer (Ambion), 2 μl 10x Dig labeling mix (Roche) and 2 μl T7 (Thermo), T3 (Roche) or SP6 (Ambion) RNA polymerase in 20 μl reactions was incubated for 2h at 37°C. To stop the reaction, 1 μl TURBO DNase I (Ambion) was added and the mix was incubated at 37°C for 15 minutes. IVT reactions were brought to 50 μl in volume and precipitated by addition of 5 μl of 3M NaOAc pH 5.2, and 125 μl of 100% EtOH and incubation at −20 °C overnight.

### *In situ* hybridization

MeOH-dehydrated embryos were rehydrated in the following series: 75% MeOH, 50% MeOH, 25% MeOH, and 1x PBST with 5-minute washes. E7.5 embryos were permeabilized with 1μl per 4 ml ProK (Invitrogen) for 1 minute at 37°C. Specimens were then incubated in glycine, post-fixed in 0.2% glutaraldehyde/4% paraformaldehyde and hybridized overnight at 65 °C in solution containing 50% formamide, 5X SSC, 1% SDS, 0.5 mg/ml tRNA (Gibco), 0.5 mg/ml Heparin (Sigma), and 100 ng/ml antisense riboprobe. Post-hybridization, embryos were washed in solution containing 50% formamide, 4X SSC and 1% SDS at 65°C, followed by washes in solution containing 500 mM NaCl, 10 mM Tris 7.5 and 0.1% Tween-20. Embryos were incubated in 100 µg/ml RNase A (Sigma) for 30 minutes at 37 °C. The embryos were then washed in MABTL made with 100 mM Maleic Acid (Sigma) pH 7.5, 150 mM NaCl, 0.1% Tween-20, 2 mM Levamisole (Sigma), and blocked in 10% sheep serum (Sigma) and 2% Boehringer Mannheim Blocking Reagent (Roche) in MABTL. Alkaline phosphatase-conjugated anti-dioxigenin FAB (Roche) fragments were then added to a solution of 1% sheep serum and 2% Boehringer Mannheim Blocking Reagent in MABTL to allow for detection of the probe. Embryos were then washed at least 10 times in MABTL to remove non-specific antibody binding. The next day, the embryos were washed in NTMTL made with 100 mM NaCl, 100 mM Tris, pH 9.5, 50 mM MgCl2, 0.1% Tween-20 and 2 mM levamisole, and then placed into color reaction with BM purple (Roche) between 4 hours and overnight at room temperature. When the color reaction was completed, the embryos were washed in PBST and post-fixed in 4% PFA. The embryos then went through washes of glycerol gradient and stored in 80% glycerol at 4°C for easier handling during imaging.

### Transcriptional Profiling of Early Embryos

Embryos used for bulk transcriptome profiling were dissected at E7.5 in 1x PBS, pooled with their littermates and dissociated with TrypLE (Fisher) reagent for 5 minutes at 37°C with shaking at 1,400 rpm in a Fisher Thermomixer followed by inactivation by addition of 10% FBS. Cells were spun down at 800 x g for 5 minutes and re-suspended in 10% FBS with a Near-IR dead dye (Life Technologies) for 30 minutes prior to sorting in order to assess cell viability. 1000 live cells were sorted directly into cell lysis buffer from either the tdTomato or YFP channel for biological replicates of wild type and mutant cells respectively. cDNA libraries were generated using SMARTer Ultra Low RNA Kit for Illumina sequencing (Clontech) and sequencing libraries were constructed using Nextera XT DNA Library Preparation Kit (Illumina). Quality control was performed both after cDNA synthesis and library preparation using Bioanalyzer (Agilent). RNA-seq Libraries were sequenced on the HiSeq2500 in the University of Chicago Functional Genomics Facility.

### Bulk RNA-seq Data Analysis

FASTQ files were aligned to the mm9 *mus musculus* genome using TopHat running standard parameters. *FeatureCounts* in *SubRead* package was used to generate read counts from the aligned bam files and subsequently analyzed using *edgeR* for differential expression. Significantly differentially expressed genes were identified using a Benjamini-Hochberg (89) corrected statistical thresholds of FDR <= 0.10. Differentially expressed genes were also used to identify associated gene ontology (GO) terms using Panther classification system (90).

### Drop-seq

#### Microfluidic Co-encapsulation of Cells and Barcoded Beads

For **Drop-seq**, 2 mL of cells at 100,000 cells/mL in PBS-BSA was loaded in a 3 mL syringe (BD, #309657). A 125 μm Drop-seq microfluidic device was used for droplet generation (31). DNA barcoded beads (ChemGenes, Macosko-2011-10(V+)) were washed, filtered, and suspended in Drop-seq lysis buffer, at 120,000 beads/mL and kept in suspension under constant stirring using a magnetic tumble stirrer and flea magnet (V&P Scientific, VP 710 Series, VP 782N-3-150). Beads and cells were co-flowed into the device, each at 4 mL/hr, along with a surfactant-oil mix (BioRad, #1864006) at 16 mL/hr that was loaded into a 10 mL syringe (BD, #302995) and used as the outer carrier oil phase. Reverse emulsions droplets were generated at ∼2000 drops/sec and collected in two batches of 15 minutes each in 50 mL tubes (Genesee Scientific, #28-106). After collection, the standard Drop-seq protocol for bead recovery, washing, and reverse transcription was followed (31). After washes and DNaseI treatment as per Drop-seq protocol (31), cDNA amplification was performed on 75,000 RNA-DNA barcode bead conjugates in a 96-well plate (Genesee Scientific, #24-302) loaded at 5000 beads per well, for a total of 15-35 wells and amplified for 14 PCR cycles using template switching (31). Post-PCR cleanup was performed by removing the STAMPs (Single Transcriptome Attached to Micro-Particles) (31) and pooling the supernatant from the wells together into a single 1.7 mL tube (Genesee Scientific, #22-281LR) along with 0.6X Ampure XP beads (Beckman Coulter, #A63880). After adding Ampure beads to the PCR product, the tube was incubated at room temperature for 2 minutes on a thermomixer (Eppendorf Thermomixer C, #5382000023) set to 1250 rpm, and for another 2 minutes on bench for stationary incubation. Next, the tube was placed on a magnet, and 4X 80% ethanol washes were performed with 1 mL ethanol added in each wash. cDNA was eluted in 150 μL of water and the concentration and library size were measured using Qubit 3 fluorometer (Thermo Fisher) and BioAnalyzer High Sensitivity Chip (Agilent, #5067-4626). 450 pg of the cDNA library was used in Nextera Library prep, instead of 650 pg as suggested in the Drop-seq protocol (31) to obtain Nextera libraries between 300 – 600 bp.

#### Sequencing

Sample libraries were loaded at ∼1.5 pM concentration and sequenced on an Illumina NextSeq 500 using the NextSeq 75 cycle v3 kits for paired-end sequencing. 20 bp were sequenced for Read 1, 60 bp for Read 2 using Custom Read 1 primer, GCCTGTCCGCGGAAGCAGTGGTATCAACGCAGAGTAC (31), according to manufacturer’s instructions. Illumina PhiX Control v3 Library was added at 5% of the total loading concentration for all sequencing runs. Each sample was sequenced on 1 or 2 lanes to a total of ∼350 – 600 million reads.

#### RNA-Seq Data Processing

Each sequencing run produced paired-end reads, with one pair representing the 12 bp cell barcode and 8 bp unique molecular identifier (UMI), and the second pair representing a 60 bp mRNA fragment (31). We utilized a pipeline (91) based on Snakemake (92) that processed raw sequencing data to produce an expression matrix corresponding to the UMI of each gene in each cell (github.com/aselewa/dropseq_pipeline). Briefly, the pipeline initially obtained a report of read quality by running *FastQC*. Next, it creates a whitelist of cell barcodes using *umi_tools* (93) *(version 0.5.3)*, based on the expected cell number per run (set to 7500 cells for this study). The cell barcode and UMI are extracted from read 1, and added to the name string of read 2 which represents the sequenced cDNA. This FASTQ file contains only the cell barcodes found in the whitelist. Finally, the protocol trims the ends of reads to remove polyA sequences and adaptors using *cutadapt* (94) *(version 1.15*). The tagged and trimmed FASTQ file is aligned to the mouse reference genome (version GRCm38/mm10) using the *STAR* (95) aligner (version 2.5.3). We used *featureCounts* (96) (version 1.6.0) to assign each aligned read to a gene on the genome. Gene annotations were based on GENCODE mV15 (97). We used the *count* function from *umi_tools* (93) to create a count matrix representing the frequency of each feature in the BAM file.

Further data processing and analyses were performed using the software tool *Seurat* (Version3) (41). Samples were imported separately and only cell barcodes with at least 1000 and less than 6000 UMIs were retained. In addition, cells with a high proportion of UMIs mapping to mitochondrial genes were removed (<5%). Data were normalized by scaling the total UMI count of each cell to 10,000 followed by log transformation (*NormalizeData*). For subsequent analyses, we selected the 2000 most variable features using *FindVariableFeatures* with default options. The datasets were combined by first identifying integration anchors (*FindIntegrationAnchors*) and then integrated (*IntegrateData*) using default settings and the first 50 PCs for both procedures. In addition to the two wildtype and two mutant samples we included a mixed sample in this run. While we were not able to assign a genotype to cells from this sample, we reasoned that the combined processing will minimize technical variation and help detection of cell types by increasing the total number of analyzed cells. This sample was used for initial integration only. Before dimensionality reduction, data were scaled using *ScaleData.* We used *RunPCA* to calculate the first 70 PCs. Only the first 45 appeared to represent significant signal and were used for further analyses. We used Uniform Manifold Approximation and Projection (UMAP) (98) applying *RunUMAP* with default settings to further reduce dimensionality. In order to cluster cells we generated the neighborhood graph using *FindNeighbors* with n.neighbors set to 20. Data were clustered using *FindClusters* with the resolution parameter set to 1.2. We found that this resolution was optimal in yielding a high number of clusters while maintaining biological interpretability. Marker genes for each cluster were identified using *FindAllMarkers* with default settings and clusters were annotated based on known marker genes and comparisons with published data (40). Based on these annotations, we subset the dataset into embryonic-derived mesoderm. We recalculated UMAP and the neighborhood graph on the subset for better visual representation and improved clustering. As before, we obtained best clustering results with resolution parameter 1.2. However, similar resolution led only to marginally different results. Marker gene and cell type identification was performed as described above. We assessed the manually identified cell types by comparison to a single cell atlas of mouse gastrulation and early embryogenesis (40). To this end we obtained the processed data generated in Pijuan-Sala (40). We generated pseudo-bulk data for both datasets by combining the UMI counts for each gene per cluster. Data were normalized by scaling each cluster to 10,000 followed by log transformation. To assess similarity between our clusters and the published clusters we obtained the pair-wise correlation between all clusters (Fig. S5).

To assess differences between wildtype and mutant cells we calculated the proportion of cells in a given cluster to the total number of cells represented by that genotype. Proportions were plotted as bar graphs. In addition, we verified that these differences were observed for both replicates. These analyses allowed us to establish differences in cellular proportions between wildtype and mutant samples in multiple clusters. However, as a change in proportion of one cluster necessarily affects the proportion of cells in other clusters, these observations alone are not sufficient to establish the directionality and primary cell type of change.

We used the data integration procedure described in Stuart et al. (41) to correct for technical differences between samples. However, this procedure could potentially ‘overcorrect’ the data and therefore minimizing biological differences between the two groups. To assess the potential magnitude of this effect we performed the same analyses without correction. As expected, these analyses showed some differences to the corrected analyses and the difference between the genotypes were more profound. However, cluster annotations were similar and thus the resulting interpretation remained unchanged (Fig. S6). Importantly, in some instances the integrated data allowed for the unambiguous identification of expected cell types (e.g pSHF) while we were unable to achieve the same resolution in the uncorrected dataset.

### Migration Assay in Chick Embryo

Chick embryos were isolated at HH stage 2/3 and cultured according to the technique described in Chapman et al., 2001 (99). DiI and DiO (ThermoFisher Scientific) were applied to the lateral aspect of the primitive streak using a Femotjet pressure injector and Injectman micromanipulator (Eppendorf) as described in Bressan et. al 2013 (81). Following 16 hrs of incubation embryos were photographed on a Lecia M165 FC fluorescent stereo microscope using a Hamamatsu Orca-flash 4.0 camera. Anterior-Posterior spread was quantified using ImageJ software (v2.0.0). Briefly, the center-of-mass for all fluorescent particles present within the embryonic area pellucida was calculated and this position was normalized to the posterior extent of the embryonic midline. For Hh inhibition studies, Cyclopamine (Sigma-Aldrich) was dissolved in 45% cyclodextrin (Sigma-Aldrich) Pannet/Compton’s saline at concentrations described in the text. 500μl of Cyclopamine solution was then added to each embryo. For FGF rescue experiments, Heparin-agarose beads (sigmaaldrich) were soaked for 1hr at room temperature in either 1mg/ml BSA or FGF4 (R&D). Beads were then implanted between the epiblast and hypoblast of HH 2/3 embryos as depicted in Figure 6 just prior to treatment with 25μM Cyclopamine.

---

**Fig. S1.**
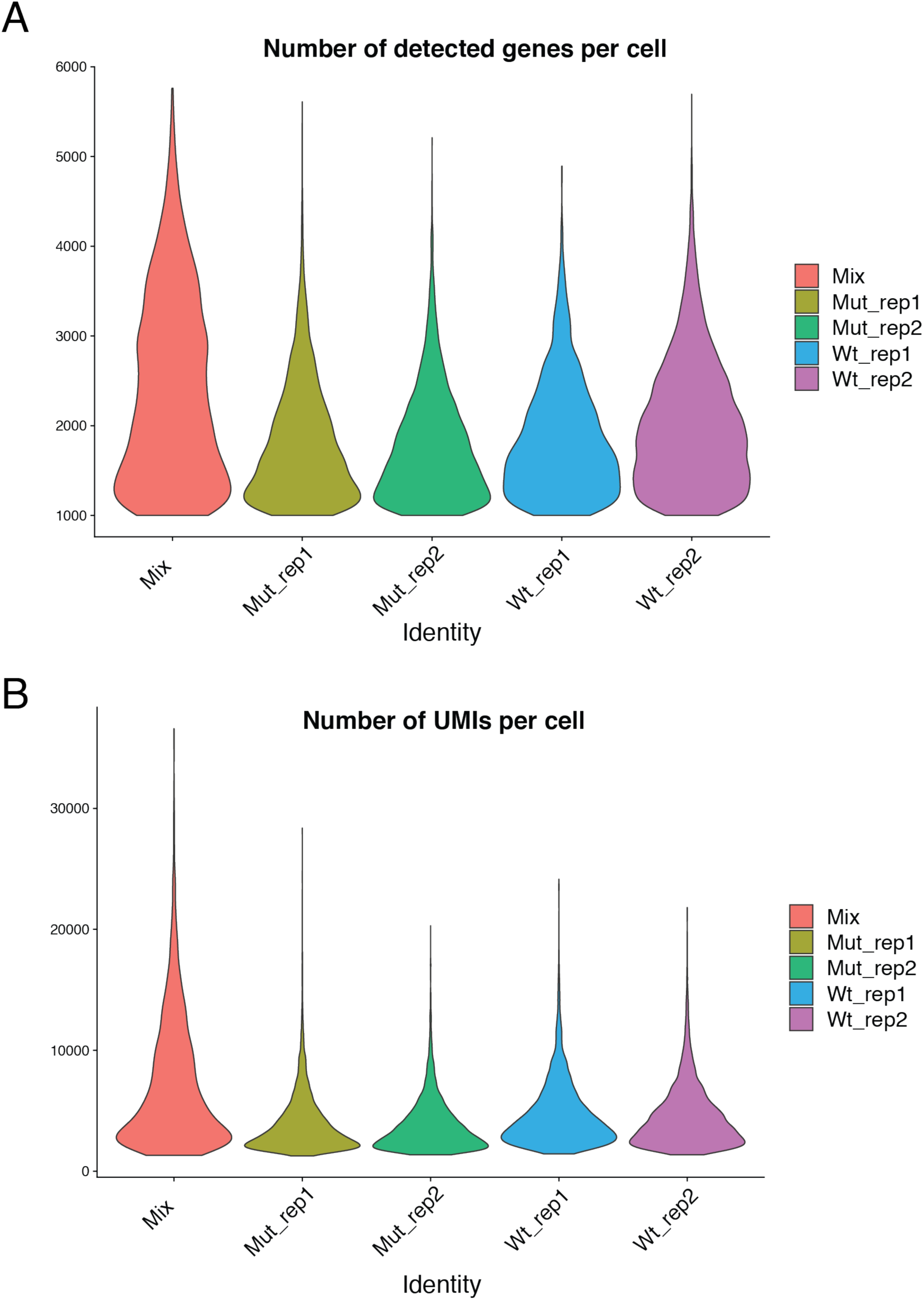
Consistent gene and UMI counts are observed between Drop-seq replicates. (*A*) Violin plots for numbers of genes detected between all samples. (*B*) Violin plots for numbers of Unique Molecular Identifiers (UMIs) detected between all cell replicates.

**Fig. S2.**
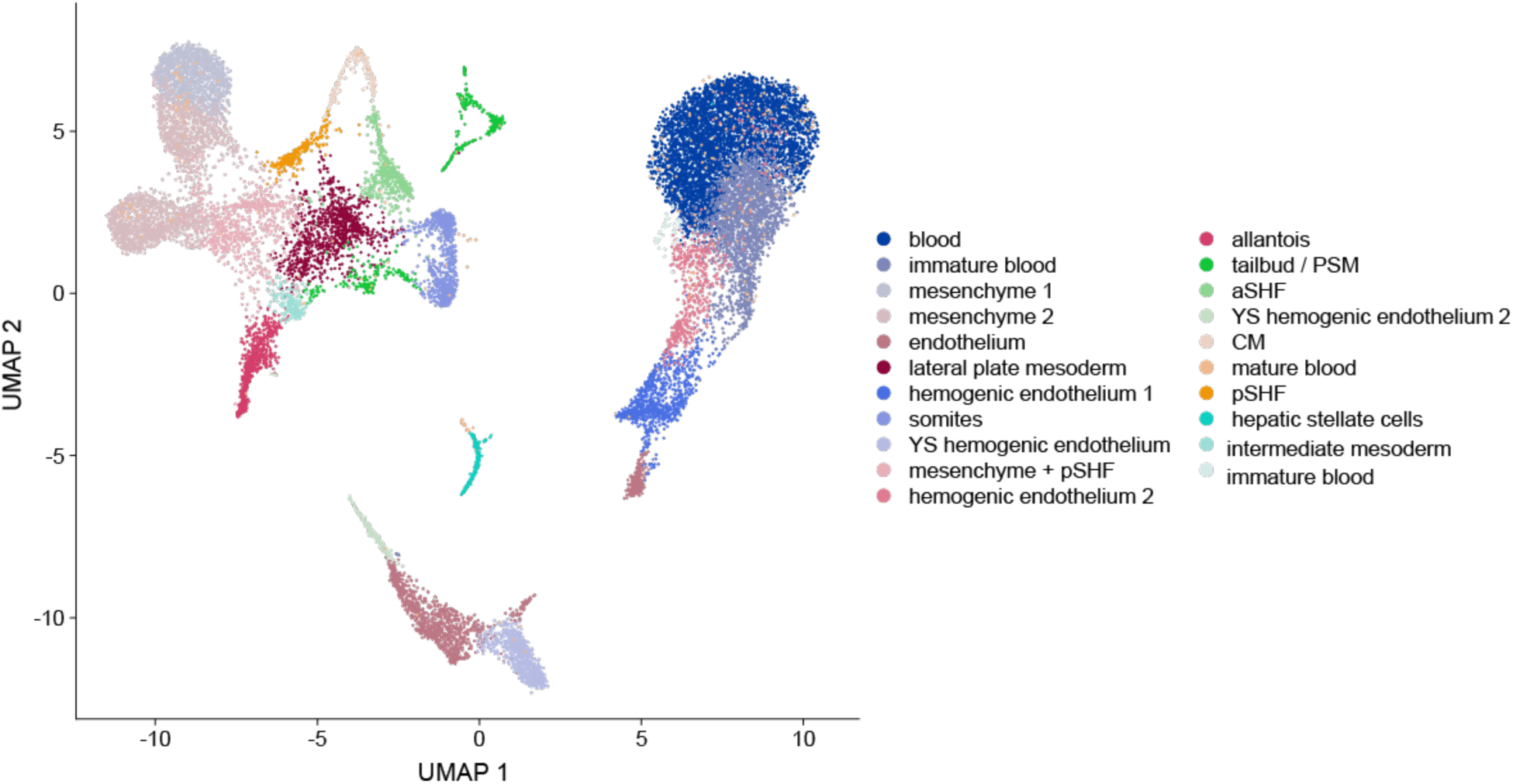
Drop-seq Assay detects all Mesp1-derived lineages from Mesp1Cre-sorted cells. UMAP for all cell types detected in Drop-seq assay for Meps1^Cre^-marked cells at E8.25.

**Fig. S3.**
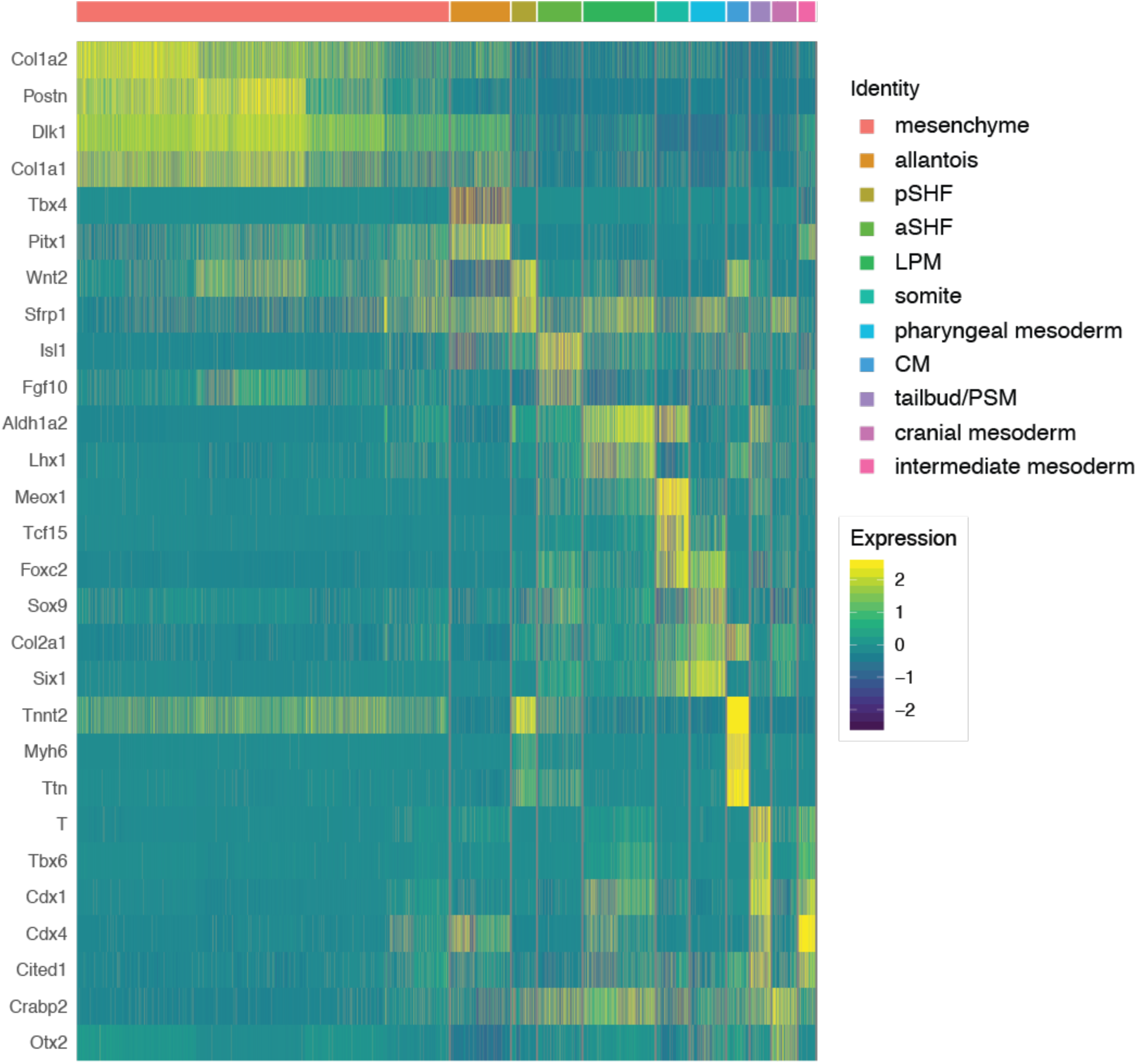
Re-clustering of embryonic mesoderm identifies all mesodermal sub-lineages by canonical marker expression. Heatmap for the expression of canonical lineage markers in sub-clustered embryonic mesoderm.

**Fig. S4.**
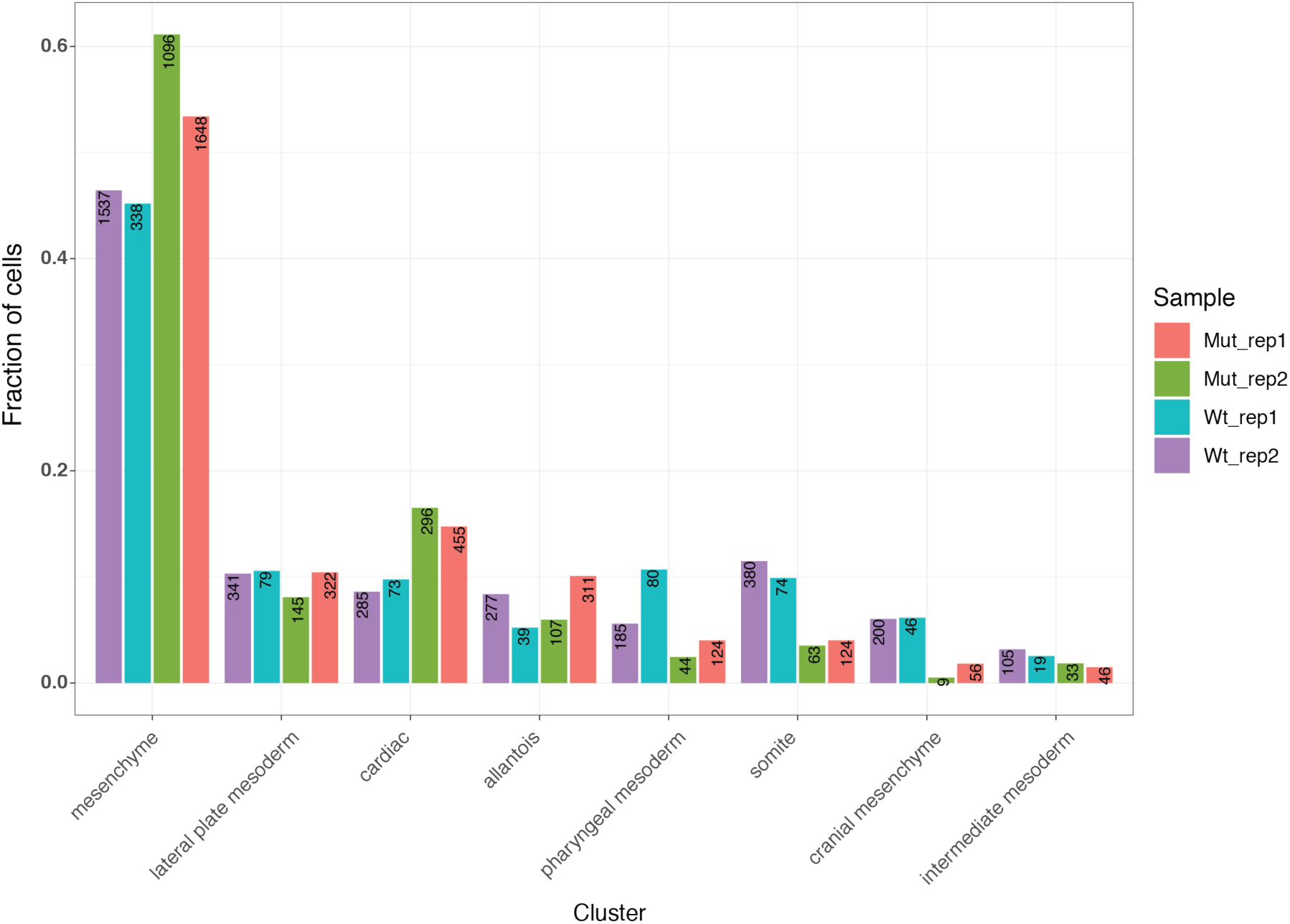
Mesoderm cell distribution is highly consistent between replicates. Column graphs representing the proportional cell contribution to each mesoderm lineage from individual replicates.

**Fig. S5.**
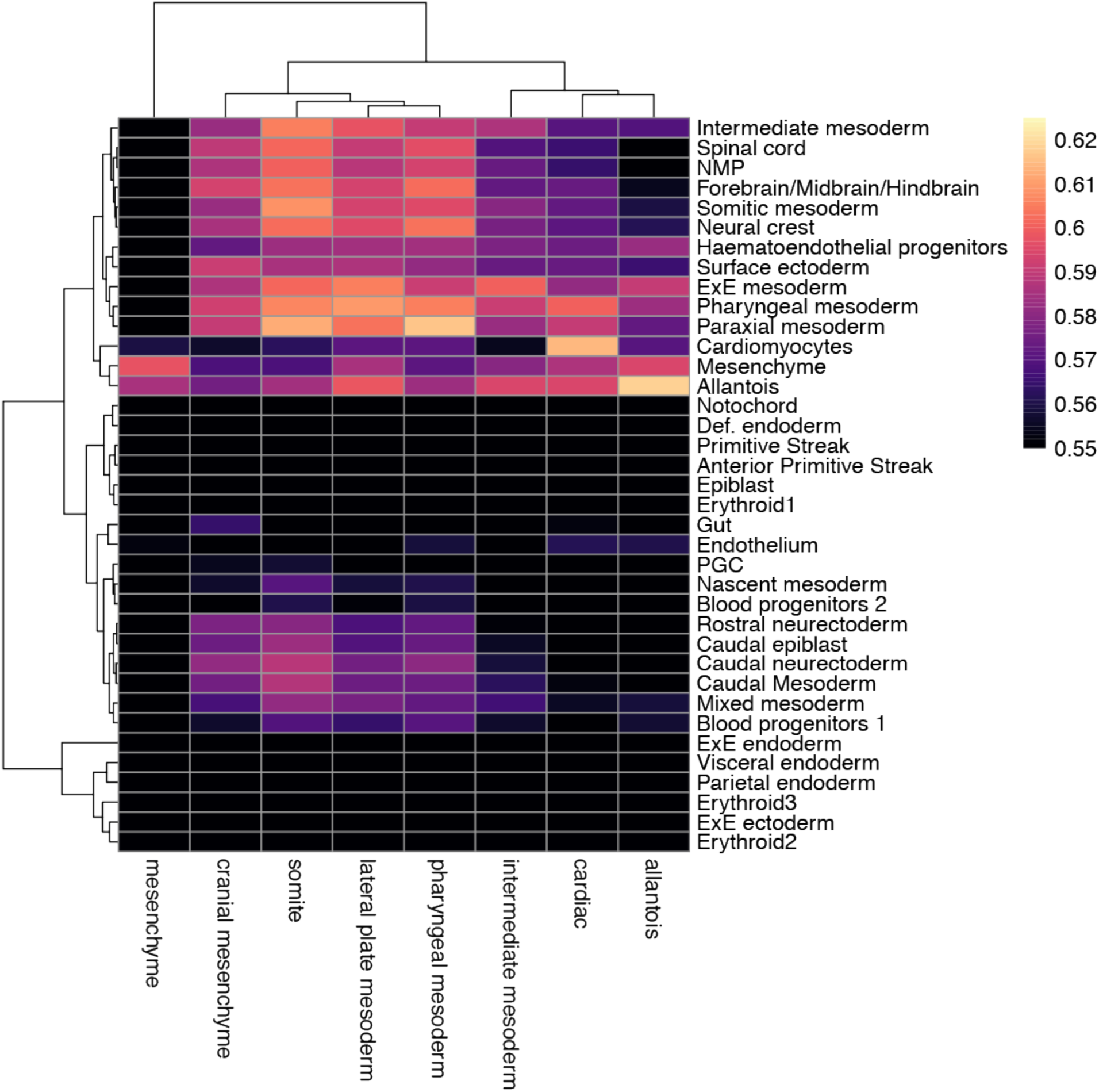
Pseudo-bulk correlation analysis between embryonic mesoderm and extant scRNA-seq datasets. Correlation matrix comparing averaged (pseudo-bulk) RNA-seq signal in each assigned embryonic mesoderm cluster with a reference whole-embryo scRNA-seq database.

**Fig. S6.**
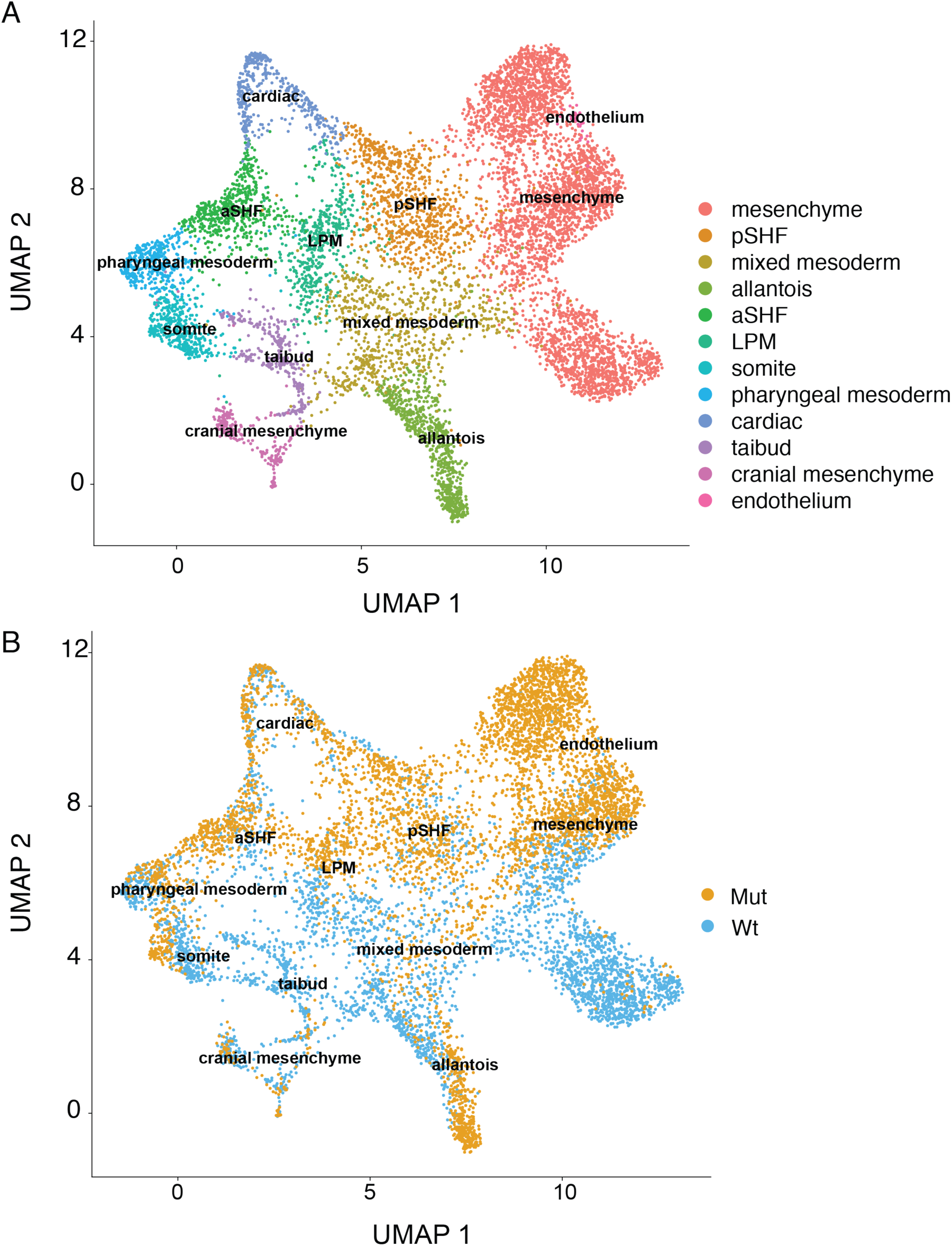
UMAP plots with non-batch-corrected scRNA-seq data. (*A* and *B*) UMAP projections demonstrating cluster assignment (*A*) and genotype (*B*).

**Fig. S7:**
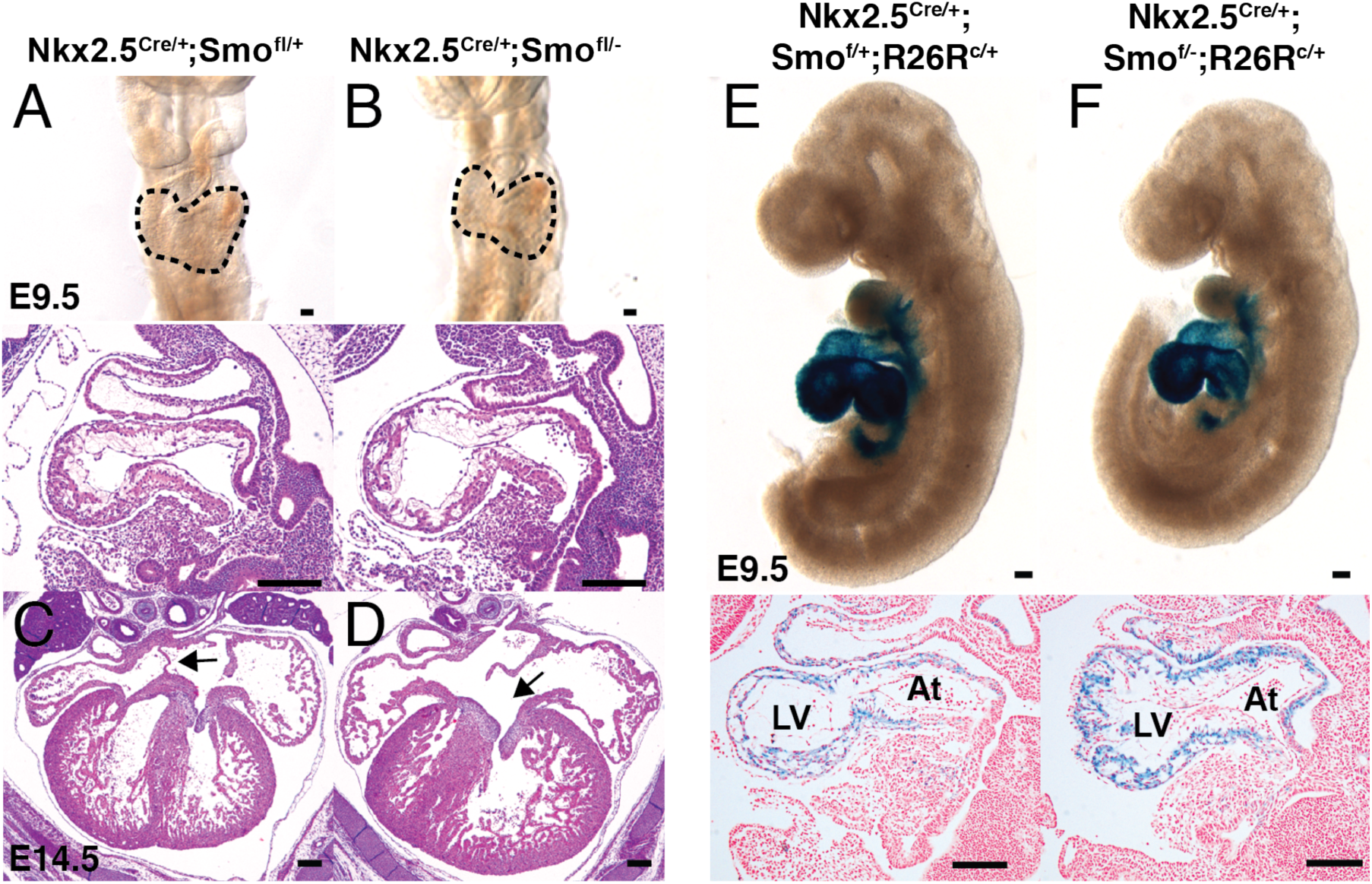
Conditional removal of Smo from early specified cardiac precursors does not cause early cardiac defects. (*A* and *B*) Shows frontal whole-mount (top panels) and sagittal histology (bottom panels) views of conditional *Smo* deletions from specified cardiac precursors in Nkx2.5^Cre/+^Smo^f/−^ E9.5 (*B*) and Nkx2.5^Cre/+^Smo^f/+^ controls (*A*). (*C* and *D*) show four-chambered-view of histology slices of E14.5 Nkx2.5^Cre/+^Smo^f/+^ and *Nkx2.5*^Cre/+^;*Smo*^f/−^ embryos; arrow points to the atrial septum (*D*). (*E* and *F*) Shows sagittal whole-mount (top panels) and sagittal histology (bottom panels) views of Nkx2.5^Cre^ fate maps in Smo^f/+^ (*E*) and Smo^f/−^ (*F*) backgrounds. Blue X-gal stain marks cells that have expressed Nkx2.5 during development and their derivatives. (Size bars = 200 µm) (LV = Left Ventricle, At = Atrium).

**Table S1.** Fate map quantification for Hh-receiving cells during gastrulation shows sparse and stochastic contribution to early cardiac structures. Values indicate numbers of marked clones in each cardiac structure derived from Gli1^CreERT2/+^;R26R^c/c^ fate maps induced by tamoxifen administration at E6.5 and E7.5 in development. Key: IFT (Inflow tract), AT (common atrium), AVC (Atrioventricular canal), LV (Left Ventricle), RV (Right Ventricle, OFT (Outflow tract).

|  | Litter 1 |  |  | Litter 2 |  |  | Litter 3 |  |  |
| --- | --- | --- | --- | --- | --- | --- | --- | --- | --- |
|  | #1 | #2 | #3 | #1 | #2 | #3 | #1 | #2 | #3 |
| **IFT** | 5 | 5 | 4 | 5 | 5 | 4 | 5 | 5 | 4 |
| **AT** | 4 | 3 | 1 | 4 | 3 | 1 | 4 | 3 | 1 |
| **AVC** | 1 | 2 | 0 | 1 | 2 | 0 | 1 | 2 | 0 |
| **LV** | 1 | 2 | 1 | 1 | 2 | 1 | 1 | 2 | 1 |
| **RV** | 2 | 3 | 1 | 2 | 3 | 1 | 2 | 3 | 1 |
| **OFT** | 3 | 4 | 3 | 3 | 4 | 3 | 3 | 4 | 3 |

